# Nuclease-NTPase systems use shared molecular features to control bacterial anti-phage defense

**DOI:** 10.1101/2025.07.10.664194

**Authors:** Adelyn E. Ragucci, Sadie P. Antine, Ethan M. Leviss, Sarah E. Mooney, Jasmine M. Garcia, Lena Shyrokova, Vasili Hauryliuk, Amy S.Y. Lee, Philip J. Kranzusch

## Abstract

Bacteria encode an enormous diversity of defense systems including restriction-modification and CRISPR-Cas that cleave nucleic acid to protect against phage infection. Bioinformatic analyses demonstrate many recently identified anti-phage defense operons are comprised of a predicted nuclease and an accessory NTPase protein, suggesting additional classes of nucleic acid targeting systems remain to be understood. Here we develop large-scale comparative cell biology and biochemical approaches to analyze 16 nuclease-NTPase systems and define shared features that control anti-phage defense. Purification, biochemical characterization, and *in vitro* reconstitution of nucleic acid targeting for each system demonstrate protein–protein complex formation is a universal feature of nuclease-NTPase systems and explain patterns of phage targeting and susceptibility. We show that some nuclease-NTPase systems use highly degenerate recognition site preferences to enable exceptionally broad nucleic acid degradation. Our results uncover shared principles of anti-phage defense system function and provide a foundation to explain the widespread role of nuclease-NTPase systems in bacterial immunity.

## Introduction

Bacteria encode hundreds of defense systems that sense and restrict bacteriophage (phage) infection^1–8^. Each bacterial genome is estimated to encode on average 5 defense systems^9^, with additional systems present in plasmids, mobile DNAs, and in phages as competition elements^10–12^. Most anti-phage defense systems are multi-gene operons, revealing a vast network of thousands of proteins dedicated to restricting phage replication^5–8,10,11,13^. Especially prevalent forms of anti-phage defense include restriction modification (RM) and CRISPR-Cas systems, present in 79% and 37% of sequenced bacterial genomes, respectively^1–4,7^. Both RM and CRISPR-Cas systems function to recognize and cleave phage DNA and RNA, demonstrating the importance of nucleic acid degradation in bacterial immunity.

Bioinformatic analyses of recently identified defense systems reveal >10% of anti-phage defense operons are predicted to encode a nuclease effector and an accessory protein with NTPase- and/or helicase-domain homology^5– 8,10,11,13–17^. These nuclease-NTPase systems lack methyltransferase or Cas proteins and support the existence of additional classes of nucleic acid-targeting systems beyond RM and CRISPR-Cas immunity. Two key examples of nuclease-NTPase immunity are the Gabija and Hachiman anti-phage defense systems. Gabija, named after the Lithuanian spirit of fire, is a two gene operon that encodes an OLD nuclease-domain containing protein (GajA) and an NTPase protein with homology to superfamily 1 helicases (GajB)^5^. Gabija is encoded in >13% of sequenced bacterial genomes and provides protection against diverse DNA phages^5,7,18^. Recent biochemical and structural studies on Gabija anti-phage defense revealed that GajA and GajB proteins interact to form a large 4:4 assembly that targets phage DNA^18–21^. Hachiman, named after a Japanese deity of war, is a similarly organized two-gene operon that encodes a PD-(D/E)XK nuclease-domain containing protein (HamA) and an NTPase protein with homology to superfamily 2 helicases (HamB)^5^. Hachiman is encoded in >5% of sequenced bacterial genomes and also provides protection against diverse DNA phages^5,7,22,23^. The Hachiman proteins HamA and HamB interact, but in contrast to Gabija form a minimal 1:1 assembly that promiscuously degrades DNA substrates *in vitro*^22^. These results demonstrate that nuclease-NTPase systems can function through a variety of biochemical and structural mechanisms and suggest that analysis of individual operons provides a limited ability to explain how nuclease-NTPase systems control anti-phage defense.

Here we develop parallel cellular and biochemical analyses of 16 diverse nuclease-NTPase systems to define general rules that control anti-phage defense. We demonstrate that nuclease-NTPase operons are broadly defensive against DNA phages and create a pairwise matrix of 319 defense system–phage combinations that reveal patterns in nuclease-NTPase targeting and phage susceptibility. Reconstituting nuclease-NTPase system function *in vitro*, we demonstrate that these systems form multimeric protein–protein complexes required for target DNA cleavage and discover that promiscuous DNA degradation exhibited by the PD-(D/E)XK nucleases in AbpAB, Hachiman, and PD-T4-8 is a result of degenerate cleavage site motifs. Our study explains how defense systems target nucleic acid during phage infection and begins to define general principles that control nuclease-NTPase system function in bacterial anti-phage defense.

## Results

### Nuclease-NTPase systems broadly defend against phage infection

To define principles that control nuclease-NTPase anti-phage defense, we analyzed nuclease-NTPase system diversity and selected representative operons for experimental studies (Figure 1a). Nuclease-NTPase systems comprise some of the most prevalent anti-phage defense operons in bacterial immunity including Gabija, Mokosh, and Hachiman^5,7,10,13^. Recent experimental and bioinformatic analyses of bacterial defense systems have uncovered >20 additional nuclease-NTPase systems that protect cells from phage infection^5,6,11,13^. Analysis of system prevalence in DefenseFinder^7^ and PADLOC^8^ demonstrates that nuclease-NTPase systems together comprise >10% of all recently described bacterial anti-phage defenses and are present in ∼40% of sequenced bacterial genomes (Figure 1a and discussed below). Nuclease-NTPase systems predominantly occur as one to three gene operons that encode a nuclease effector protein associated with a partnering NTPase, helicase, and/or a domain of unknown function protein^5,6,11,13– 17^.

**Figure 1.**
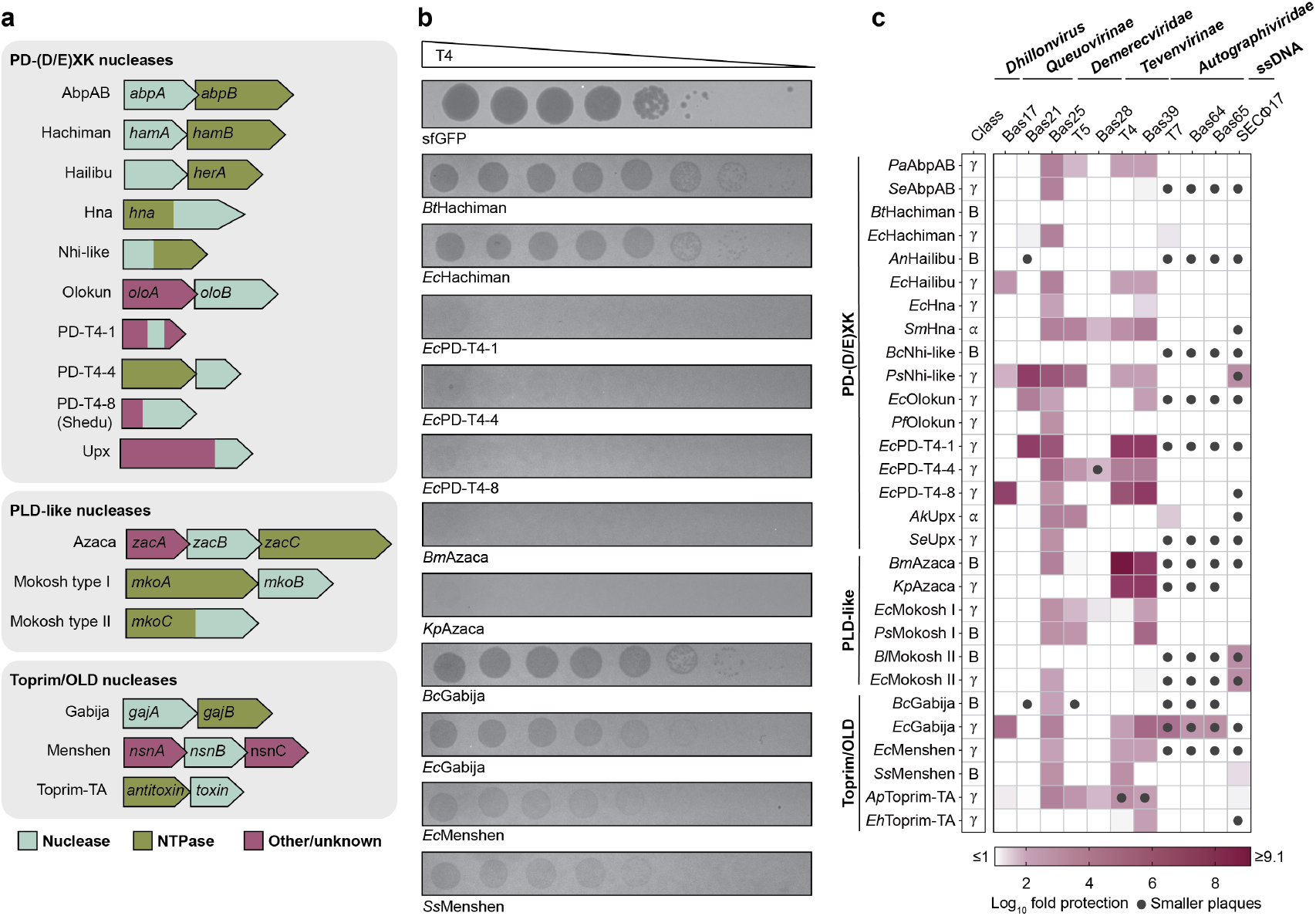
Nuclease-NTPase systems broadly defend against DNA phage infection. **a**, Schematics of individual genes in selected nuclease-NTPase operons. Systems are grouped based on nuclease type with nuclease proteins or domains depicted in blue, NTPase proteins or domains in green, and other domains or domains of unknown function in pink. **b**, Representative plaque assays for bacteria expressing nuclease-NTPase operons challenged with phage T4. *Ec*PD-T4-1, *Ec*PD-T4-4, *Ec*PD-T4-8, *Bm*Azaca, *Kp*Azaca, *Ec*Menshen, and *Ss*Menshen provide defense against T4. **c**, Heatmap displaying the log10 fold protection conferred by nuclease-NTPase operons against a panel of phages. In the leftmost column, operons which native hosts are *Gammaproteobacteria* are indicated with a η, *Alphaproteobacteria* with a, and *Bacilli* are indicated with a B. Data in **b–c** are representative of at least three independent experiments.

Analysis of the nuclease effector proteins in nuclease-NTPase systems reveals three major groups: PD-(D/E)XK (InterPro IPR011604), phospholipase D (PLD)-like (InterPro IPR025202), and Toprim/OLD nucleases (InterPro IPR006171) (Figure 1a). The PD-(D/E)XK nuclease family members exhibit low overall sequence similarity but share a common core aβββaβ fold characteristic of endonucleases encoded in restriction-modification systems^24^. Conserved PD-(D/E)XK active site residues are critical for coordinating metal-dependent nucleophilic hydrolysis of phosphodiester bonds in a reaction that can be adapted for cleavage of both RNA and DNA target substrates^24–26^. As PD-(D/E)XK enzymes represent some of the most common forms of nuclease proteins in nuclease-NTPase systems, we selected 10 systems from this nuclease class including AbpAB, Hachiman, Hailibu, Hna, Nhi-like, Olokun, PD-T4-1, PD-T4-4, PD-T4-8, and Upx (Figure 1a). PLD enzymes include prokaryotic and eukaryotic proteins involved in phospholipid metabolism, including cardiolipin synthases^27^ and the piRNA pathway protein, Zucchini^28^. These enzymes use an invariant H(X)K(X4)D active-site motif to catalyze a two-step mechanism that can cleave DNA substrates through creation of a phosphohistidine intermediate^26,27^. We selected 2 nuclease-NTPase systems with PLD-like enzymes: Azaca and Mokosh (Figure 1a). Finally, Toprim (topoisomerase-primase) domain containing proteins harbor two conserved motifs (a conserved glutamate and two aspartates DXD) and catalyze a concerted metal-dependent reaction that induces double stranded DNA (dsDNA) breaks^26,29^. OLD (overcoming lysogeny defect) proteins, first described in the phage P2, contain an N-terminal ATP-binding cassette (ABC)-ATPase and a C-terminal Toprim domain^30^. It has been suggested that OLD nucleases cleave DNA via a canonical two-metal DNA cleavage mechanism^26,31,32^. We selected 3 nuclease-NTPase systems with Toprim/OLD nucleases: Gabija, Menshen, and Toprim-TA^5,13,17^ (Figure 1a).

In addition to the nuclease effector, most nuclease-NTPase systems encode an accessory NTPase protein with predicted homology to Superfamily 1 (SF1) or Superfamily 2 (SF2) helicases^33,34^. SF1 and SF2 helicases share high sequence homology, including >9 amino acid motifs, and overall structural similarity, but differ primarily in conservation of motifs III and IV and contacts made by these regions^35^. The bacterial helicase UvrD, known for its role in DNA recombination and repair, defines the family of SF1 helicases^33,36^. SF2 helicases consist of many sub-families, including RNA unwinding Ski-2-like and the RIG-I-like translocases^34^. Of the 16 nuclease-NTPase systems we selected for analysis, the NTPase accessory proteins in Gabija, Mokosh, and Nhi-like are SF1 helicases and the accessory proteins in AbpAB, Azaca, Hachiman, Hna, and Toprim-TA are SF2 helicases (Figure 1a). A defining feature of all SF1 and SF2 helicases is the presence of a Walker A motif (also known as motif I, AxxGxGK(T/S)) that forms the phosphate binding loop required for interaction with magnesium and ATP, and a Walker B motif (also known as motif II, DEXX) that coordinates magnesium and directs NTP hydrolysis^33^. In addition to classical helicase enzymes, Walker A and Walker B motifs are also associated with proteins that do not exhibit conventional nucleic acid unwinding, suggesting that not all partnering accessory proteins in nuclease-NTPase systems may be involved in nucleic acid interactions^34^. The NTPase domains in nuclease-NTPase system proteins also share predicted homology with P-loop NTPases which typically hydrolyze ATP or GTP as primary substrates^37^. The accessory proteins in AbpAB, Azaca, Gabija, Hachiman, Hailibu, Hna, Menshen, Mokosh, Nhi-like, PD-T4-4, and Toprim-TA all have predicted homology to P-loop NTPases and contain conserved Walker A motifs. Across nuclease-NTPase systems, the NTPase accessory protein is typically larger than the nuclease effector. Within the operons selected for our analysis, the gene order across diverse systems does not appear to be conserved with an approximately equal distribution of which gene is encoded first (Figure 1a).

We assayed each of the 16 selected nuclease-NTPase systems for the ability to defend against phage infection. For each nuclease-NTPase system, we aimed to select two representative operon sequences, one from *E. coli* or the nearest available proteobacteria species, and one from *Bacillus* or a firmicute bacteria that exhibited ∼20–60% aminoacid identity to the proteobacteria system across the nuclease and helicase protein coding regions (Figure 1a, Table S1). In most cases, we included the sequence of the nuclease-NTPase operon which was verified to be phage-defensive upon its discovery, and for the PD-T4 systems we chose to only study the operons from *E. coli*. We cloned the native operon sequences (including gene overlaps and intergenic regions) into an arabinose inducible vector for expression in an *E. coli* strain engineered to remove 9 endogenous cryptic prophages (*E. coli* BW25113 Δ9)^38^. Using a titration series of arabinose induction, we selected an induction concentration of 0.02% to ensure expression levels with minimal observable impact of operon toxicity on bacterial growth (Extended data figure 1). We challenged *E. coli* expressing each of the 29 nuclease-NTPase operons with a panel of 10 diverse dsDNA coliphages from the BASEL collection^39^ and one ssDNA phage SECФ17, thus creating a matrix of 319 pairwise interactions that reveals general principles of nuclease-NTPase immunity (Figure 1b,c).

First, nuclease-NTPase anti-phage defense is phage specific with different patterns of protection evident against distinct phages as well as viral families. All nuclease-NTPase operons tested were able to defend against at least one phage (Figure 1b,c). Some phages like phage Bas25 are particularly susceptible and were potently restricted by most nuclease-NTPase operons, but other phages like Bas21 and Bas28 are generally resistant to nuclease-NTPase anti-phage defense. In some cases, including T7-like phages from the family *Autographiviridae*, the presence of a nuclease-NTPase operon had relatively little effect on viral titer but notably reduced plaque size (Figure 1c, Extended data figure 2). Additionally, nuclease-NTPase operons originating from proteobacteria genomes were overall more defensive against *E. coli* phages than firmicute homologs, suggesting that these operons are specifically adapted for function in proteobacteria species or may respond to coliphage-specific cues as has been observed with other anti-phage defense systems^40–42^ (Figure 1c). Second, some nuclease-NTPase operons appear to function as generalist systems, broadly restricting replication of diverse DNA phages, while other operons appear specialized and potently restrict a narrow range of phages. The most broadly defensive operons in our panel were *Ec*Gabija and *Ps*Nhi-like, each exhibiting >20-fold defense against 7 of the 11 phages tested (Figure 1c). Additional generalist antiphage defense operons included *Pa*AbpAB, *Sm*Hna, and *Ap-* Toprim-TA that broadly defend against many phages but exhibit an overall weaker defense profile with modest reduction in phage output titer. Specialized nuclease-NTPase systems with selective phage restriction include Azaca that provides >1,000,000-fold protection against phage T4 and Mokosh type II that only defends against the ssDNA phage SECФ17 (Figure 1b,c). Additionally, select individual operons potently restrict phages that are otherwise generally resistant to nuclease-NTPase defense with *Ec*PD-T4-8 capable of restricting phage Bas17 and *Ec*PD-T4-1 and *Ps*Nhi-like capable of restricting phage Bas21 (Figure 1c). Interestingly, in our analysis of nuclease-NTPase systems we did not note patterns of defense specific to nuclease protein family or helicase type. Together, these results provide a matrix of defense operon–phage interactions that reveal defense profiles across diverse nuclease-NTPase systems and suggest that nuclease-NTPase systems utilize distinct mechanisms not determined by specific protein classes.

### Complex formation is a widespread feature of nuclease-NTPase systems

To begin defining mechanisms controlling nuclease-NTPase anti-phage defense, we next purified representative proteins from each system for biochemical analyses. We cloned variants of each multi-gene operon with a 6×His-SUMO2 tag on the N-terminus of the first protein in the operon or a 6×His tag on the C-terminus of the final operon protein and over-expressed the constructs in *E. coli* for purification. We successfully purified recombinant protein from 14 of 16 representative nuclease-NTPase systems excluding DUF4297-HerA (recently re-named Hailibu)^43–45^ and Mokosh Type I (Extended data figure 3). Remarkably, purification of individual tagged proteins in each case resulted in co-isolation of all associated defense system component proteins supporting widespread formation of large protein assemblies (Extended data figure 3a). These results agree with previous analyses of Gabija and Hachiman systems^19–23,46^ and demonstrate that higher-order complex formation between nuclease proteins and accessory NTPase subunits is a defining feature broadly conserved in nuclease-NTPase immunity.

Previous structural analyses revealed that Gabija forms a giant 496 kDa supramolecular complex composed of a 4:4 GajA–GajB nuclease-NTPase assembly^19–21,46^. In contrast, the Hachiman nuclease-NTPase system forms a simple 1:1 HamA–HamB assembly that is 160 kDa in size^22,23^. To define how mechanisms of complex formation vary across diverse nuclease-NTPase systems, we used mass photometry (MP) to analyze complex size for each purified system^47^ (Figure 2). MP analysis of *Bc*Gabija determined an experimental mass of ∼490 kDa, accurately matching the GajA–GajB assembly known to be required for Gabija antiphage defense^19–21,46^ (Figure 2f). Nuclease-NTPase system complexes in our panel exhibited a broad range of sizes *in vitro* ranging from minimal ∼140 kDa systems like *Pa*AbpAB (∼142 kDa), *Ec*Mokosh II (∼138 kDa), and *Se*Upx (∼141 kDa) to larger higher-order assemblies like *Pf*Olokun (∼278 kDa) and *Ec*PD-T4-8 (∼193 kDa) (Figure 2). To define potential complex subunit stoichiometries, we used the experimental MP data as restraints to guide AlphaFold3 structural modeling of each nuclease-NTPase system^48^ (Figure 2, Extended data figure 4). Starting first with minimal assemblies, MP analysis was consistent with monomeric structural models for Mokosh type II (132 kDa) and Upx (143 kDa), and 1:1 complexes for Hachiman (1:1 HamA–HamB) and AbpAB (1:1 AbpA–AbpB) (Figure 2a,b,d,e). The AbpA–AbpB predicted model closely resembles a recent cryo-EM structure of the HamA–HamB complex, further supporting shared homology between these two systems^22,23^. Notably, although the nuclease subunit structures of AbpA and HamA superimpose well, AbpA contains an additional C-terminal DUF4297 Cap4 nuclease domain that is predicted to extend outside the HamA–HamB interface and form unique contacts with the NTPase subunit AbpB^23,49^ (Figure 2a,b). Interestingly, the AbpA N-terminal nuclease domain resembles HamA but lacks conserved active site residues suggesting that enzymatic function is carried out exclusively by the AbpA DUF4297 Cap4 domain. MP analysis of Hna was consistent with monomeric (97 kDa) and dimeric (194 kDa) states in solution, and AlphaFold3 generated high-confidence models for each state (Figure 2c). In both Hna predicted models, the PD-(D/E)XK nuclease domain resides at the outer edge of the complex in a near identical placement of the same nuclease domain in the predicted structure of Upx (Figure 2e).

**Figure 2.**
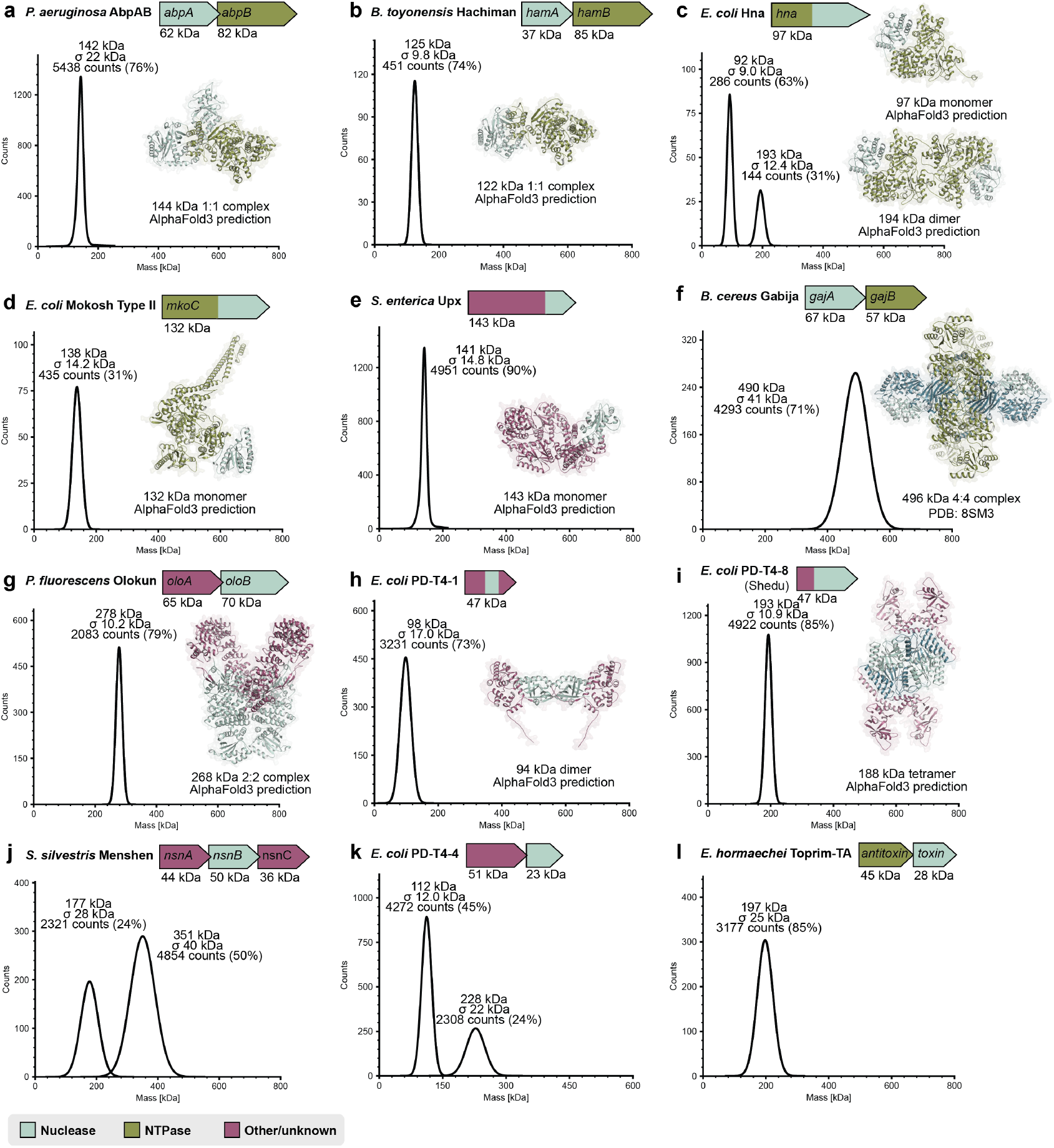
Complex formation is a widespread feature of nuclease-NTPase systems. **a–l**, Mass photometry analysis of purified nuclease-NTPase complexes. Peaks are labeled with molecular weight, standard deviation of the Gaussian fit, and particle counts. Nuclease-NTPase operons are shown above and models of experimentally determined complex structures or AlphaFold3 structural predictions are shown to the right of each graph. **a–e**, Minimal nuclease-NTPase complexes. *Pa*AbpAB (**a**) and *Bt*Hachiman (**b**) form 1:1 complexes. *Ec*Hna (**c**) exists as both a monomer and a dimer in solution. *Ec*Mokosh II (**d**) and *Se*Upx (**e**) are monomeric. **f–i**, Higher order nuclease-NTPase complexes. *Bc*Gabija (**f**) forms a 4:4 supramolecular complex. *Pf*Olokun (**g**) forms a 2:2 complex. *Ec*PD-T4-1 (**h**) and *Ec*PD-T4-8 (**i**) are dimeric and tetrameric, respectively. **j–l**, *Ss*Menshen (**j**), *Ec*PD-T4-4 (**k**), and *Eh*Toprim-TA (**l**) MP graphs have multiple peaks and/or high standard deviations which prevent reliable determination of stoichiometric state. Data shown are representative of at least 2 independent experiments.

For the larger nuclease-NTPase higher-order assemblies, combined MP and AlphaFold3 analyses allowed identification of a single best structural model for most systems. These analyses support that Olokun forms a 2:2 OloA–OloB complex with the adaptin-like OloA subunit interacting with the N-terminal helical domain of the nuclease OloB (Figure 2g). The OloB PD(D/E)XK nuclease domains are predicted to reside adjacent to one another in a configuration potentially allowing formation of doublestranded breaks. PD-T4-1 and PD-T4-8 systems are predicted to form dimeric and tetrameric complexes, respectively, with the PD-(D/E)XK nuclease domains forming the complex core (Figure 2h,i). In further support of our analyses, PD-T4-8 is equivalent to the Shedu anti-phage defense system which was recently demonstrated to form a tetrameric complex nearly identical to our predicted model^50,51^. MP analysis of Menshen, PD-T4-4, and Toprim-TA systems demonstrates formation of higher-order complexes, but MP data contained multiple peaks with high standard deviation preventing reliable stoichiometry prediction (Figure 2j,k,l). Technical limitations with complex stability during MP analysis prevented guided prediction of Azaca and Nhi-like systems structure models. Our analyses define experimentally supported structure models for diverse nuclease-NTPase systems and provide a foundation for future structural determination efforts.

### Nuclease-NTPase systems have distinct DNA cleavage patterns

We next developed biochemical assays to define how nuclease-NTPase systems target and degrade nucleic acid. Leveraging our purification of 14 representative nuclease-NTPase systems, we tested each system for the ability to degrade a panel of 11 target nucleic acid substrates including host *E. coli* chromosomal DNA, closed circular plasmid DNA, and phage genomic DNA purified from the virions of 7 dsDNA and 2 ssDNA phages (Figure 3a–c). Notably, from our target DNA panel the genomic DNA of phages T4, lambda, and SPO1 are known to be hyper-modified with nucleobase modifications^52^. Three major phenotypes emerged from our comparative analysis of *in vitro* cleavage specificity. First, the purified nuclease-NTPase complexes of *Pa*AbpAB, *Bt*Hachiman, and *Ec*PD-T4-8 were highly active and promiscuously cleaved all DNA targets *in vitro* (Figure 3c, Extended data figure 5). Each of these three systems encode PD-(D/E)XK nuclease domains demonstrating this nuclease type is capable of broadly targeting and cleaving methylated DNA and DNA with diverse nucleobase modifications. Although *Pa*AbpAB, *Bt*Hachiman, and *Ec*PD-T4-8 completely degraded every tested DNA substrate *in vitro*, no toxicity was observed upon recombinant protein overexpression of these systems in *E. coli* indicating that regulatory mechanisms not recapitulated in our minimal biochemical system exist in cells to limit nucleic acid degradation in the absence of infection. Second, many nuclease-NTPase complexes were active *in vitro* but exhibited specific targeting and degradation of only select phage genomic DNAs (Figure 3c, Extended data figure 5). *Ps*Nhi-like and *Pf*Olokun robustly degraded phage T7, SECФ17, and SPO1 genomic DNA but were unable to target *E. coli* genomic and plasmid DNA suggesting that these systems may use specific structural or sequence features to discriminate DNA targets. The *in vitro* DNA cleavage patterns of *Ps*Nhi-like and *Pf*Olokun only partially correlate with the ability of these systems to defend against phage infection *in vivo*, suggesting that target DNA specificity may broaden upon full activation *in vivo* and that some phages may avoid system activation or use mechanisms of immune evasion to prevent restriction (Figure 1c, Figure 3c, Extended data figure 5). The nuclease-NTPase operons *Ec*PD-T4-1 and *Ec*Mokosh II only cleaved ssDNA targets in our panel and exhibited no cleavage for any dsDNA target (Figure 3c, Extended data figure 5). It is possible that these systems may recognize long ssDNAs to discriminate host DNA from viral replication intermediates present during infection. *Bm*Azaca exhibited a clear preference for phage T4 dsDNA degradation, agreeing with strong *in vivo* defense of Azaca operons against phage T4 and related T-even phages (Figure 1c, Figure 3c, Extended data figure 2,5). Phage T4 genomic DNA is modified with bulky glucosyl-hydroxymethylcytosine modifications providing a possible molecular cue for Azaca target recognition^53^. Finally, some nuclease-NTPase operons including *Ec*Hna, *Ec*PD-T4-4, *Se*Upx, *Bc*Gabija, *Ss*Menshen, and *Eh*Toprim-TA were inactive under all conditions tested (Figure 3c, Extended data figure 5). It is possible these nuclease-NTPase systems may require a specific viral cue for activation or target nucleic acids like RNA or tRNA not included in our panel.

**Figure 3.**
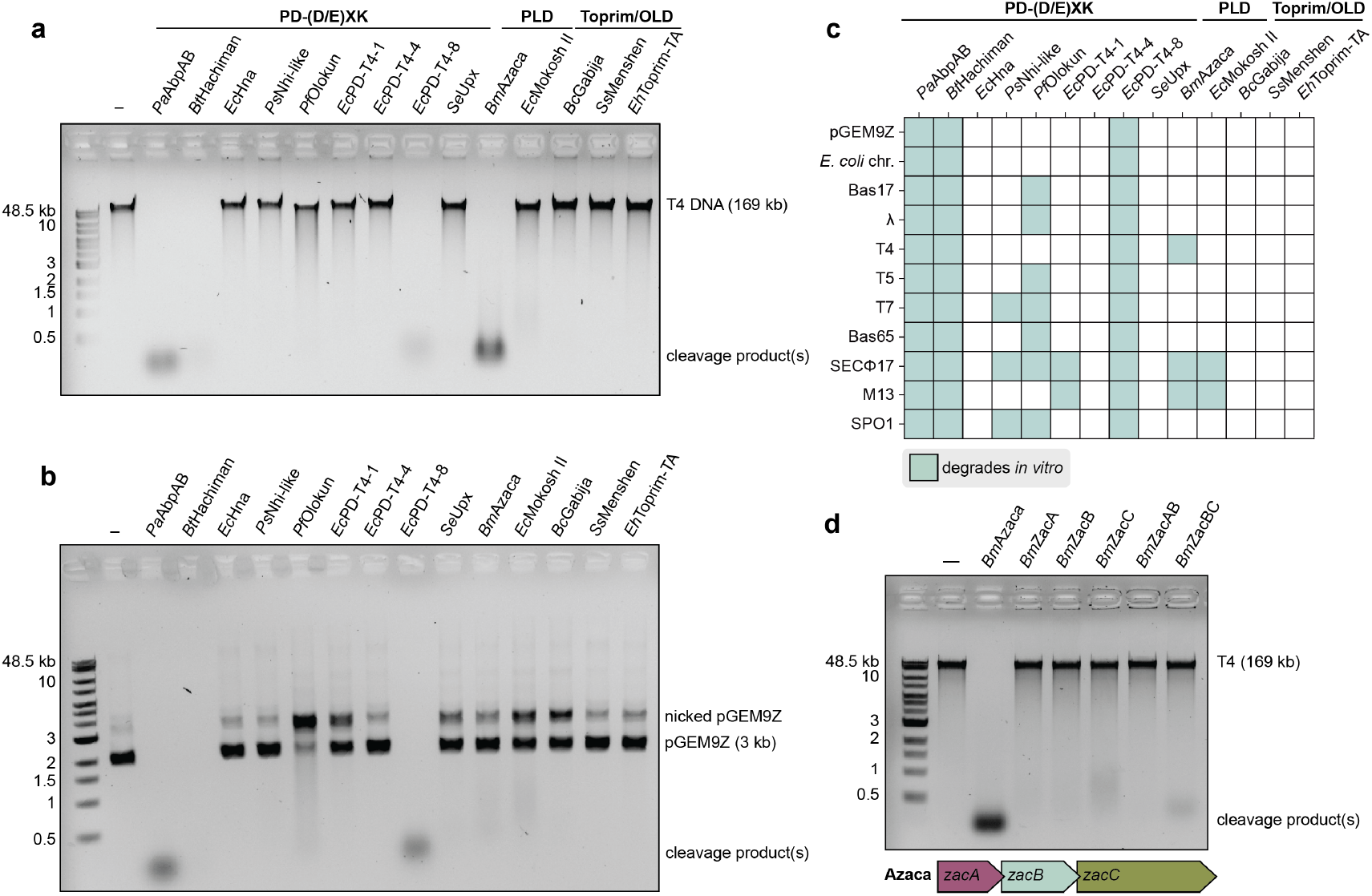
Nuclease-NTPase systems have distinct DNA cleavage patterns. **a**, DNA cleavage and agarose gel analysis of 14 nuclease-NTPase systems incubated with phage T4 genomic DNA. *Pa*AbpAB, *Bt*Hachiman, *Ec*PD-T4-8, and *Bm*Azaza degrade phage T4 genomic DNA. **b**, DNA cleavage and agarose gel analysis of nuclease-NTPase systems incubated with plasmid DNA. *Pa*AbpAB, *Bt*Hachiman, and *Ec*PD-T4-8 degrade plasmid DNA substrate while *Pf*Olokun, *Ec*PD-T4-1, *Ec*Mokosh II, and *Bc*Gabija induce reduced substrate mobility consistent with plasmid nicking activity. **c**, Summary of nuclease-NTPase system in vitro cleavage of host and phage DNA with nuclease-NTPase systems on the x-axis and phages on the y-axis. A filled in square indicates nucleic acid cleavage. **d**, DNA cleavage and agarose gel analysis of the ability of the individual or pairwise subunits of Azaca to degrade phage T4 DNA. Only the *Bm*Azaca complex consisting of all three proteins (ZacABC) cleaves phage T4 DNA. Data in **a–d** are representative of at least 3 independent experiments.

We next examined active site residues, conserved motifs, and individual subunit requirements for nuclease-NTPase systems. In each case, disruption of conserved nuclease active site residues led to complete loss of dsDNA cleavage activity (Extended data figure 6a–i). Mutations to the conserved DEK nuclease active site residues of the DUF4297 domain of *Pa*AbpAB (*Pa*AbpA*B) and the DUF1837 domain in *Bt*Hachiman (*Bt*HamA*B) abolished dsDNA cleavage but these mutant variants retained the ability to nick plasmid DNA as evidenced by conversion of target plasmid to a lower mobility DNA species (Extended data figure 6a,d). For most systems, disruption of nuclease active site residues had no impact on nuclease-NTPase complex formation, but as an exception we observed that mutations to the *Pf*Olokun OloB nuclease active site caused loss of OloAB complex formation during purification. Mutations that disrupt the conserved NTPase Walker A and Walker B active site motifs in *Pa*AbpB, *Bt*HamB, and *Ec*MkoC had no impact on DNA cleavage activity, however disruption of the Walker A motif in *Bm*ZacC resulted in complete loss of target DNA cleavage (Extended data figure 6a,c,d,f). The Azaca operons also contain an uncharacterized ZacA protein with a DUF6361 domain, and mutations to highly conserved residues in this protein region had no impact on complex formation or target DNA cleavage (Extended data figure 6c). In Gabija anti-phage defense, the nuclease subunit GajA is alone sufficient to direct DNA cleavage *in vitro*^18,19^. We therefore purified individual nuclease domains in isolation for *Pa*AbpAB, *Bt*Hachiman, and *Bm*Azaca. In contrast to Gabija, we observed that in each case DNA degradation required the NTPase subunits AbpB, HamB, and ZacC (Figure 3d, Extended data figure 6b,e). Furthermore, we observed that in the case of *Bm*Azaca, all three Azaca system proteins (ZacA, ZacB, and ZacC) are necessary for DNA cleavage (Figure 3d). Together, these results define nucleic acid target specificity for diverse nuclease-NTPase systems and reveal critical roles for conserved active site residues and higher-order complex formation.

### Nuclease-NTPase systems exhibit degenerate cleavage site motifs

We next investigated the molecular basis of how nuclease-NTPase systems control DNA target site recognition. Activation of nuclease enzymes in bacterial anti-phage defense frequently requires concerted organization of multiple protomers and nuclease domain clustering^1,3^. To begin to define the impact of protein concentration on DNA target site recognition, we tested activity of each nuclease-NTPase system *in vitro* over a range of protein concentrations from 10 nm to 1 µM. *Pa*AbpAB, *Bm*Azaca, *Ec*Mokosh II, and *Ec*PD-T4-1 each exhibited robust nuclease activity at all concentrations tested suggesting that these nuclease-NTPase systems are capable of functioning as individual or minimally organized complexes and do not require significant clustering (Figure 4a–c; Extended data figure 7b,e). In contrast, *in vitro* reactions with *Bt*Hachiman, *Ps*Nhi-like, *Pf*Olokun, and *Ec*PD-T4-8 nuclease-NTPase operons were only active at higher 1 µM concentrations indicating that these systems may require molecular crowding for activation or that under these minimal *in vitro* conditions molecular crowding is necessary to achieve activation in the absence of phage-associated cues (Figure 4a–c; Extended data figure 7a,c,d,f).

**Figure 4.**
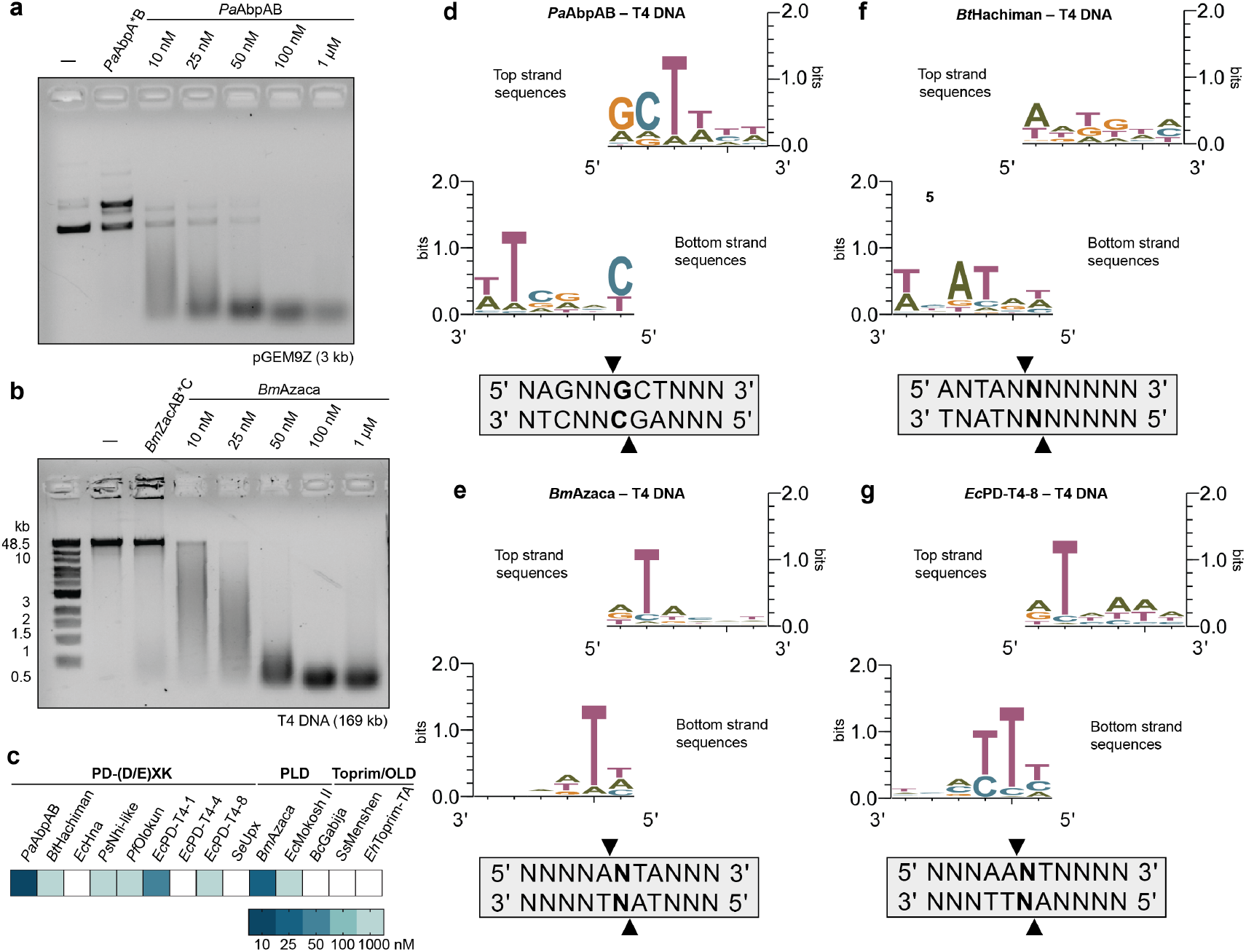
Nuclease-NTPase systems exhibit degenerate cleavage site motifs. **a**, DNA cleavage and agarose gel analysis of the ability of *Pa*AbpAB to cleave plasmid DNA at protein concentrations from 10 nM to 1 µM demonstrates *Pa*AbpAB is an active nuclease at concentrations as low as 10 nM. **b**, DNA cleavage and agarose gel analysis of the ability of *Bm*Azaca to cleave phage T4 DNA at protein concentrations from 10 nM to 1 µM demonstrates *Bm*Azaca begins to cleave at 10 nM. **c**, Summary of nuclease-NTPase system concentration dependence data. *Pa*AbpAB, *Ec*PD-T4-1, and *Bm*Azaca cleave DNA at concentrations !50 nM. *Bt*Hachiman, *Ps*Nhilike, *Pf*Olokun, *Ec*PD-T4-8, and *Ec*Mokosh II cleave DNA only at 1 µM. Empty squares indicate systems which were not active under these conditions. **d–g**, Cut-site analysis of phage T4 DNA fragments from *Pa*AbpAB (**d**), *Bm*Azaca (**e**), *Bt*Hachiman (**f**), and *Ec*PD-T4-8 (**g**) reactions using next generation sequencing reveals that these nuclease-NTPase systems use highly degenerate sequence motifs. Data in **a–c** are representative of at least 3 independent experiments.

Using sequencing analysis of DNA cleavage reactions, we defined the cleavage site motifs for *Pa*AbpAB, *Bm*Azaca, *Bt*Hachiman, and *Ec*PD-T4-8^54–56^. Deep sequencing of DNA fragments revealed that each nuclease-NTPase system uses a highly degenerate, minimized cleavage site motif to direct DNA targeting (Figure 4d–g, Extended data figure 8b–g). *Pa*AbpAB DNA target fragments exhibit a consensus cleavage site motif of 5’ N↓GCT (Figure 4d, Extended data figure 8b). *Bt*Hachiman has a highly degenerate consensus sequence with minimal to no sequence preference consistent with the indiscriminate DNA cleavage observed *in vitro* (Figure 4f, Extended data figure 8e). Both *Ec*PD-T4-8 and *Bm*Azaca DNA target fragments exhibit a consensus cleavage site of 5’ A↓NT (Figure 4e,g, Extended data figure 8g). Consistent with highly degenerate motifs, mapping of the observed cut sites demonstrated that each nuclease-NTPase system cleaved all regions throughout the plasmid and T4 genomic DNA targets. Additionally, analysis of parallel reactions performed with plasmid DNA or phage T4 genomic DNA for *Pa*AbpAB, *Bt*Hachiman, and *Ec*PD-T4-8 revealed the same cleavage site preferences for each system regardless of DNA target type (Figure 4d−g, Extended data figure 8b,e,g). Together our results define the cleavage site motifs of diverse nuclease-NTPase systems and explain how minimal sequence preferences enable a broad range of DNA target cleavage specificity.

## Discussion

Our study of 29 diverse operons enables comparative analysis of the role of nuclease-NTPase systems in bacterial anti-phage defense. Parallel *in vivo* and biochemical experiments with each nuclease-NTPase system provide an important complement to previous in-depth mechanistic characterization of individual operons^14–16,18,19,22,23,43,44,50,57^. Our data synthesize with previous results to begin to reveal general rules that control how nuclease-NTPase systems function (Figure 5). Major principles that emerge include (1) nuclease-NTPase operons can function as both general and highly selective immune systems to provide broad antiphage defense (Figure 1), (2) protein–protein complex formation is a shared nuclease-NTPase system feature that controls nuclease activity and protein stability (Figures 2 and 3), and (3) nuclease-NTPase systems appear to primarily induce nonselective nucleic acid cleavage without sequence-specific target recognition (Figures 3 and 4).

**Figure 5.**
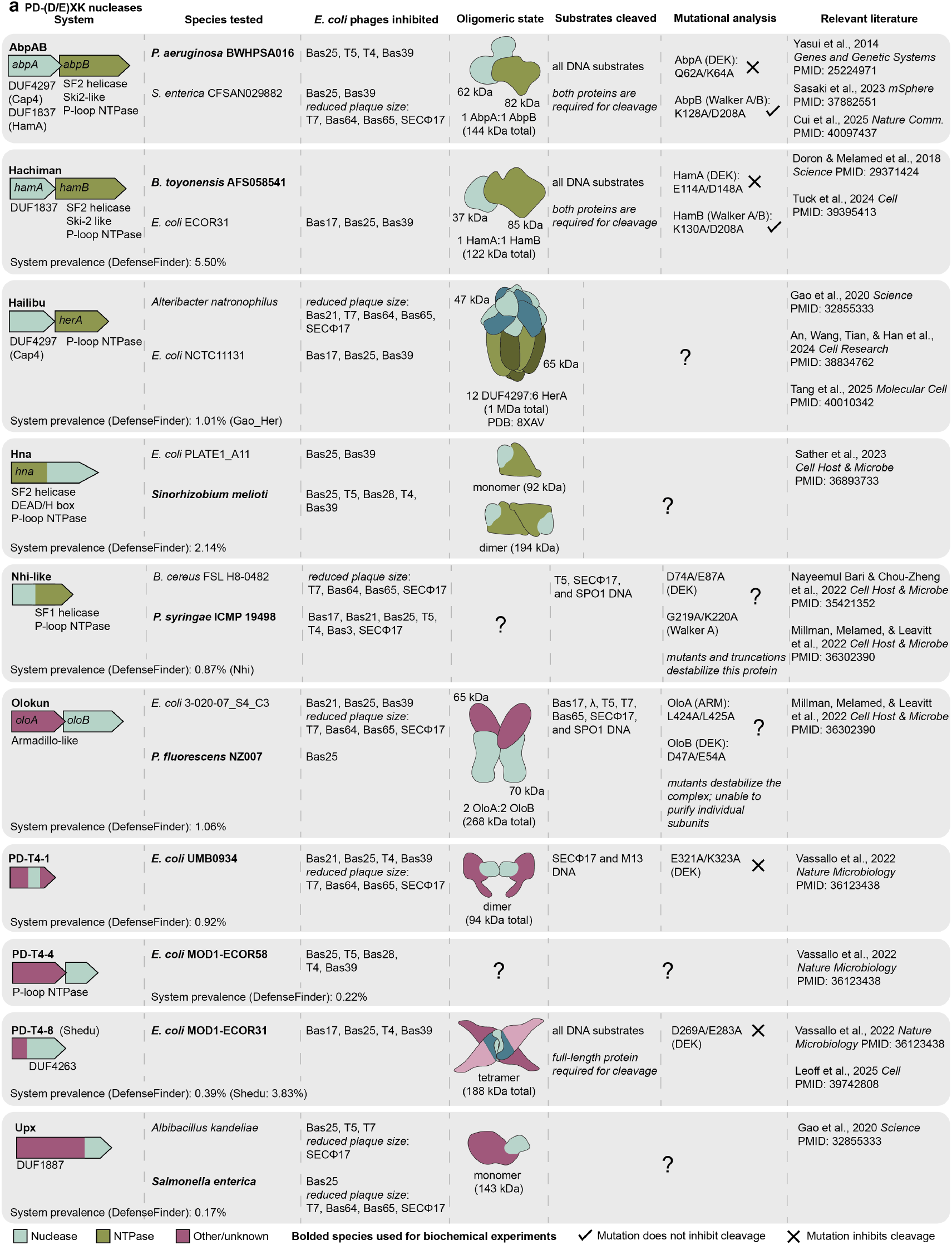
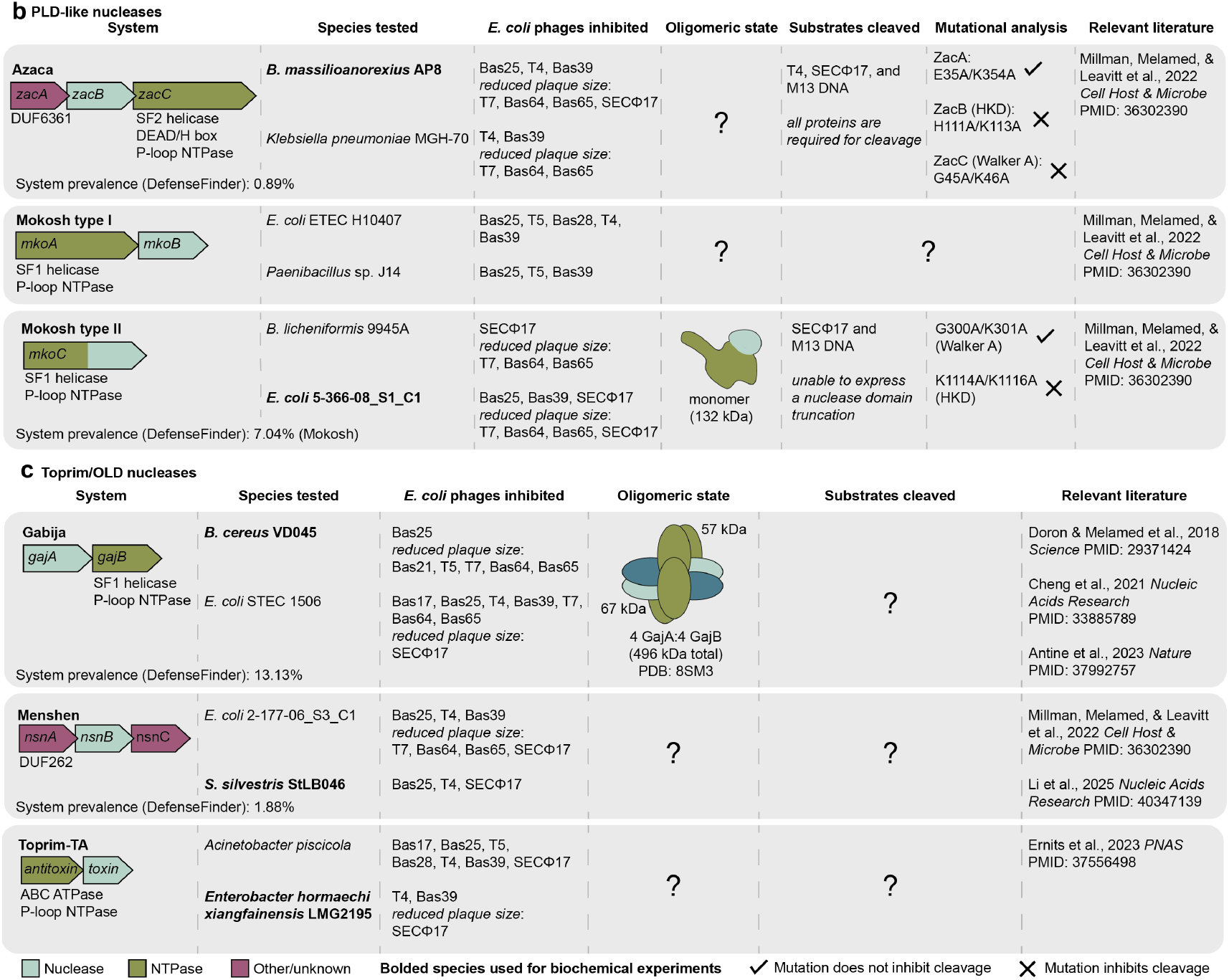
A detailed reference for nuclease-NTPase anti-phage defense systems. **a–c**, A comprehensive summary of known information for each nuclease-NTPase system included in this study. Nuclease-NTPase operons are grouped according to nuclease domain homology with PD-(D/E)XK nucleases (**a**), PLD-like nucleases (**b**), and Toprim/OLD nucleases (**c**) operons presented.

Nuclease-NTPase operons are capable of defending *E. coli* against a broad diversity of DNA phages. Screening 11 DNA phages representing 5 viral families, our results create a matrix of operon-phage interactions that allow direct comparisons between nuclease-NTPase systems (Figure 1). Some nuclease-NTPase systems exhibit exceptionally broad defense with the *Ec*Gabija and *Ps*Nhi-like operons in our screen defending against more than half of all phages tested and other operons like *Pa*AbpAB, *Sm*Hna, and *Ap*Toprim-TA exhibit defense against at least 4 diverse phages in our panel (Figure 1c, Extended data figure 2). In contrast, other nuclease-NTPase systems like Azaca and Mokosh II defend strongly but only against select subsets of phages (Figure 1c, Extended data figure 2). Our results do not support a correlation between individual nuclease or NTPase protein domains and form of anti-phage defense, suggesting that even nuclease-NTPase systems with related components can control distinct phage restriction patterns *in vivo*.

Nuclease-NTPase systems function through protein– protein complex formation and share features conserved across bacterial and animal cell antiviral immunity. We observed that the protein subunits from each characterized nuclease-NTPase operon interact to form a discrete complex, supporting that complex formation is a potentially universal feature of this form of immune defense (Figure 2). In animal immunity, antiviral signaling is typically controlled by large, oligomeric protein complexes that coordinate pathogen sensing and signal transduction^58–60^. Similarly, nuclease-NTPase systems join other well-characterized bacterial antiphage defense systems that function through supramolecular complex formation including CRISPR systems, RADAR, PARIS, and effector proteins in CBASS and Thoeris defense^3,4,49,61–65^. Nuclease-NTPase systems can form a minimal 1:1 complex (AbpAB and Hachiman) or significantly more complex higher-order assemblies (Gabija, Olokun, and PD-T4-8), suggesting that alterations to complex formation may enable evolutionary adaption to target distinct aspects of phage infection or limit viral immune evasion^19,43,61,64,66^.

Our results demonstrate that nuclease-NTPase systems can function through a diverse array of nuclease effector subunits including PD-(D/E)XK, PLD-like, and OLD nuclease domains. Mutational analyses confirmed that system activity was strictly dependent on the canonical nuclease active site residues (Extended data figure 6). PD-(D/E)XK nuclease domains include classical restriction enzymes that inhibit phage infection by catalyzing sequence-specific cleavage of conserved nucleic acid motifs^24^. In contrast, our results demonstrate that PD-(D/E)XK nucleases in nuclease-NTPase systems are highly promiscuous and cleave diverse target nucleic acids using highly degenerate sequence motifs (Figure 3 and Figure 4). In our analysis, nuclease-NTPase systems with PLD-like nuclease subunits exhibited more defined substrate specificity, and systems with OLD nuclease subunits did not exhibit cleavage activity with model nucleic acid substrates *in vitro* (Figure 3). These results suggest that some nucleases, like OLD nuclease domains, may be particularly specialized to target specific nucleic acid substrates like hairpin or branched intermediate structures only present during phage replication^67^.

Major open questions remain in understanding how nuclease-NTPase systems restrict phage replication. For nearly all nuclease-NTPase systems, how phage infection is sensed remains unknown or poorly understood^18,19,22,40^. One potential model is that complex activation and phage target recognition *in vivo* is controlled primarily by the NTPase subunit. In our biochemical experiments, nuclease subunits alone were inactive and complex formation between the nuclease and NTPase subunits was required for nucleic acid cleavage (Extended data figure 6). The Gabija nuclease-NTPase system is an exception to these results, with the *Bacillus cereus* GajA nuclease subunit known to be sufficient to degrade short, specific phage lambda sequences *in vitro*^18,19^. For nearly all tested nuclease-NTPase systems including Gabija, the NTPase subunit active site motifs responsible for NTP hydrolysis are essential for phage defense *in vivo* but are dispensable for nucleic acid degradation *in vitro*^5,6,13–15,19^ (Extended data figure 6), These results suggest that minimal nuclease-NTPase target cleavage reactions reconstituted *in vitro* likely artificially bypass normal aspects of phage sensing and complex regulation. In cells, NTPase subunits may be required for loading nuclease-NTPase complexes on relatively rare phage nucleic acid substrates that occur at lower concentrations than model nucleic acids presented *in vitro*. Alternatively, nuclease-NTPase complex assembly may also be required *in vivo* to limit aberrant nuclease activity similar to the common role of regulatory subunits that restrain nucleases in type II toxin-antitoxin systems^68^. Our results create a new foundation to test these and other hypotheses and explain further general properties that control how nuclease-NTPase operons function in bacterial anti-phage defense.

## Acknowledgements

The authors are grateful to members of the Kranzusch laboratory for helpful comments and discussion. We thank the Center for Macromolecular Interactions at Harvard Medical School and the Dana-Farber Cancer Institute Molecular Biology Core Facility. We thank Gemma Atkinson for sharing the sequence of the Toprim-TA system. The work was funded by grants to P.J.K. from the Pew Biomedical Scholars program, the Burroughs Wellcome Fund PATH program, The G. Harold and Leila Y. Mathers Charitable Foundation, The Mark Foundation for Cancer Research, the Cancer Research Institute, the Parker Institute for Cancer Immunotherapy, and the National Institutes of Health (1DP2GM146250-01), grants to A.S.Y.L. from The G. Harold and Leila Y. Mathers Charitable Foundation and the National Institutes of Health (R35GM142527), and grants to V.H. from the Knut and Alice Wallenberg Foundation (project grant 2020-0037 V.H.), the Swedish Research Council (Vetenskapsrådet) grants (2021-01146 and 2024-06059 to V.H.), and the Göran Gustafsson Foundation for Research in Natural Sciences and Medicine (the Göran Gustafsson Prize to V.H.).

J.M.G is supported by a Ford Foundation Predoctoral Fellowship and the National Institutes of Health (5TL1TR002543). This paper was typeset with the bioRxiv word template by @Chrelli: www.github.com/chrelli/bioRxiv-word-template

## Author contributions

The study was designed and conceived by A.E.R., S.P.A. and P.J.K. Phage challenge experiments were performed by A.E.R and S.E.M. Protein purification and biochemical assays were performed by A.E.R, S.P.A, E.M.L, and S.E.M. Bacterial growth assays were performed by

S.E.M and E.M.L. Mass photometry and AlphaFold3 modeling was performed by S.P.A. Sequencing analysis was performed by J.M.G and

A.S.Y.L. L.S and V.H. performed initial phage defense assays with the Toprim-TA system. Figures were prepared by A.E.R. with contributions from S.P.A and E.M.L. The manuscript was written by A.E.R. and P.J.K. All authors contributed to editing the manuscript and support the conclusions.

## Data availability statement

Sequencing data generated from this study will be made available at the NCBI Sequence Read Archive. Information required to reanalyze the sequencing data is available from the lead contact upon request. All other data are available in the manuscript or the supplementary information.

## Materials and Methods

### Bacterial growth and phage defense assays

Nuclease-NTPase operons were synthesized and cloned into an arabinose-inducible pBAD vector (GenScript)^69^. *E. coli* BW21113 Δ9 (NCTC 14365)^38^ cells were transformed with nuclease-NTPase operon plasmids and plated onto LB supplemented with 1% glucose and 100 µg mL^−1^ ampicillin. For bacterial growth assays, one colony from each transformation was picked into 3 mL LB with 1% glucose and 100 µg mL^−1^ ampicillin and grown overnight at 37°C with shaking at 230 rpm. Cells were pelleted, resuspended in PBS (137 mM NaCl, 2.7 mM KCl, 10 mM Na_2_HPO_4_, 1.8 mM KH_2_PO_4_), and serially diluted in a 96-well plate. 5 µL of each serial dilution was spotted onto LB plates supplemented with 100 µg mL^−1^ ampicillin and 0.02% L-arabinose as well as LB plates supplemented with 100 µg mL^−1^ ampicillin and 1% glucose. Plates were allowed to dry for 1 hour and incubated overnight at 37°C. For plaque assays, one colony from each transformation was picked into 5 mL LB supplemented with 100 µg mL^−1^ ampicillin and grown at 37°C for 6 hours with shaking at 230 rpm. Cells were normalized to an OD_600_ of 0.3 and mixed with SM top agar (LB with 0.1 mM MnCl_2_, 5 mM MgCl_2_, 5 mM CaCl_2_, 100 µg mL^−1^ ampicillin, 0.02% arabinose, and 0.8% agar) to total 15 mL. Plates were left to solidify for 45 mins at room temperature before 2.5 µL drops of serially diluted phage lysate were pipetted on top of the solidified agar layer containing the bacteria. Spots were allowed to dry before incubation overnight at 30°C. Fold defense was calculated as the ratio between PFU mL^−1^ for phages plated on sfGFP expressing bacteria compared to nuclease-NTPase operon expressing bacteria. When individual plates could not be counted, a faint lysis zone was counted at 10 plaques^70^. Bacterial growth and plaque assay plates were imaged on a ChemiDoc MP Imaging System. When necessary, brightness and contrast were adjusted for visualization of colonies and plaques.

### Nuclease-NTPase system selection and plasmid construction

Designated nuclease-NTPase systems were selected from the literature with the requirement of having a nuclease domain encoded in the operon and a predicted NTPase/helicase superfamily accessory protein. If applicable, the operon reported in the original discovery paper and a diverse homolog from a different bacterial species were synthesized and cloned into custom pET vectors with an N-terminal 6×His-SUMO2 fusion tag on the first protein in the operon or a C-terminal 6×His tag on the last gene in the operon (Genscript)^71^. The original host bacterial sequence was used leaving the sequence of the operon intact from the start codon of the first gene to the stop codon of the last gene. Individual genes were cloned into either pET vectors with an N-terminal 6×His-SUMO2 fusion tag on the first protein in the operon or a C-terminal 6×His tag on the last protein in the operon. Active site mutations were synthesized by Genscript or cloned using site directed mutagenesis^72^. Full operon protein expression vectors and active site mutants used for protein expression and purification are as follows: N-terminal 6×His-SUMO2 *Pa*AbpAB, N-terminal 6×His-SUMO2 *Bm*Azaca, 6×His-SUMO2-5×GS *Bc*Gabija, N-terminal 6×His-SUMO2 *Bt*Hachiman, *Ec*Hna C-terminal 6×His tag, *Ss*Menshen C-terminal 6×His tag, *Ec*Mokosh II C-terminal 6×His tag, N-terminal 6×His-SUMO2 *Ps*Nhi-like, N-terminal 6×His-SUMO2 *Pf*Olokun, *Ec*PD-T4-1 C-terminal 6×His tag, *Ec*PD-T4-4 C-terminal 6×His tag, 6×His-SUMO2-5×GS *Ec*PD-T4-8, *Eh*Toprim-TA C-terminal 6×His tag, and N-terminal 6×His-SUMO2 *Se*Upx. Individual protein expression vectors are as follows: N-terminal 6×His-SUMO2 *Pa*AbpA, N-terminal 6×His-SUMO2 *Pa*AbpB, N-terminal 6×His-SUMO2 *Bm*ZacA, N-terminal 6×His-SUMO2 *Bm*ZacB, N-terminal 6×His-SUMO2 *Bm*ZacC, N-terminal 6×His-SUMO2 *Bm*ZacAB, N-terminal 6×His-SUMO2 *Bm*ZacBC, N-terminal 6×His-SUMO2 *Bt*HamA, N-terminal 6×His-SUMO2 *Bt*HamB, *Ec*Mokosh II M1-A995 C-terminal 6×His tag, *Ec*Mokosh II L996-end C-terminal 6×His tag, N-terminal 6×His-SUMO2 *Pf*OloA, and N-terminal 6×His-SUMO2 *Pf*OloB, N-terminal 6×His-SUMO2 *Ps*Nhi-like M1–F163, N-terminal 6×His-SUMO2 *Ps*Nhi-like E164–end, N-terminal 6×His-SUMO2 *Ec*PD-T4-8 M1–D192, N-terminal 6×His-SUMO2 *Ec*PD-T4-8 K193–end.

### Protein expression and purification

Nuclease-NTPase complexes were expressed in *E. coli* as previously described (Zhou 2018 Cell PMID: 30007416). Briefly, expression vectors were transformed into BL21(DE3)-RIL (Agilent), or LOBSTR-BL21(DE3)-RIL cells (Kerafast), plated on MDG media plates (1.5% Bacto agar, 0.5% glucose, 25 mM Na_2_HPO_4_, 25 mM KH_2_PO_4_, 50 mM NH_4_Cl, 5 mM Na_2_SO_4_, 0.25% aspartic acid, 2–50 µM trace metals, 100 µg mL^−1^ ampicillin, 34 µg mL^−1^ chloramphenicol) and grown overnight at 37°C. Three to five colonies were picked to inoculate 30−60 mL overnight MDG starter cultures (37°C, 230 rpm). 1 L M9ZB expression cultures (47.8 mM Na_2_HPO_4_, 22 mM KH_2_PO_4_, 18.7 mM NH_4_Cl, 85.6 mM NaCl, 1% Cas-Amino acids, 0.5% glycerol, 2 mM MgSO_4_, 2–50 µM trace metals, 100 µg mL^−1^ ampicillin, 34 µg mL^−1^ chloramphenicol) were then inoculated with 10 mL or starting OD_600_ of 0.0475 of MDG starter cultures and then induced with 0.5 M IPTG after reaching an OD_600_ of ≥1.5 followed by overnight induction (16°C, 230 rpm).

After overnight inductions, cells were centrifuged, resuspended, and pellets were lysed by sonication in 60 mL lysis buffer (20 mM HEPES pH 7.5, 400 mM NaCl, 10% glycerol, 20 mM Imidazole, 1 mM DTT). Lysate was clarified by centrifugation and resulting supernatant was added to Ni-NTA resin (Qiagen). Resin was washed with lysis buffer, lysis buffer supplemented to 1 M NaCl, and then lysis buffer again for *Pa*AbpAB, *Bc*Gabija, *Bt*Hachiman, *Ec*Hna, *Ec*Mokosh II, *Ps*Nhi-like, *Ec*PD-T4-1, and *Se*Upx. The resin was washed with lysis buffer three times for *Bm*Azaca, *Ss*Menshen, *Pf*Olokun, *Ec*PD-T4-4, and *Eh*Toprim-TA. Samples were eluted with lysis buffer supplemented to 300 mM Imidazole. Samples were then dialyzed overnight in 14 kDa MWCO dialysis tubing (Ward’s Science) with SUMO2-cleavage by hSENP2 as previously described^71^. *Bm*Azaca was dialyzed and stored in gel filtration buffer (20 mM HEPES-KOH pH 7.5, 20 mM KCl, and 1 mM TCEP-KOH) supplemented with 10% glycerol. *Ec*Mokosh II was next purified by ion-exchange chromatography using 5 mL HiTrap Heparin HP columns (Cytiva) and eluted by a 150 mM – 1 M NaCl gradient. Fractions were pooled and further purified by size exclusion chromatography using a 16/600 Sephacryl 300 column (Cytiva) and stored in gel filtration buffer. *Bc*Gabija and *Ps*Nhi-like were purified by size exclusion chromatography using a 16/600 Sephacryl 300 column (Cytiva) and stored in gel filtration buffer. *Pa*AbpAB, *Bt*Hachiman, *Ec*Hna, *Ss*Menshen, *Pf*Olokun, *Ec*PD-T4-1, *Ec*PD-T4-4, *Ec*PD-T4-8, *Eh*Toprim-TA, and *Se*Upx were purified by size exclusion chromatography using a 16/600 Superdex 200 column (Cytiva) and stored in gel filtration buffer. Final proteins were concentrated to >10 mg mL^−1^ using a 30 kDa MWCO centrifugal filter (Millipore Sigma), aliquoted, flash frozen in liquid nitrogen, and stored at −80°C.

All catalytic active site mutants were purified by Ni-NTA, dialyzed and stored in gel filtration buffer (20 mM HEPES-KOH pH 7.5, 20 mM KCl, and 1 mM TCEP-KOH) supplemented with 10% glycerol with comparison to the WT protein/complex. Individual proteins or domains from each system were purified by Ni-NTA dialyzed and stored in gel filtration buffer (20 mM HEPES-KOH pH 7.5, 20 mM KCl, and 1 mM TCEP-KOH) supplemented with 10% glycerol with comparison to the full protein or protein complex.

### Mass photometry and structural modeling

The oligomeric state of purified proteins was determined by mass photometry using a Refeyn MP instrument (TwoMP, Refeyn) RRID:SCR_018270^47^. Purified proteins were diluted to 200–400 nM in gel filtration buffer prior to analysis. Isopropanol and MilliQ-H_2_O cleaned coverslips were loaded with gaskets and placed on the objective lens with a drop of Immersol immersion oil (Zeiss). AcquireMP software (Refeyn) was used to find focus on gel filtration buffer only, and then beta-amylase and thyroglobulin control or samples were added to the buffer 1:10 followed by immediate recording for 60 seconds. Ratiometric contrast measurements were converted to molecular weight measurements using DiscoverMP software (Refeyn). Automatic Gaussian curve fitting was performed on all oligomeric states.

The AlphaFold3 server was used to predict the structure of nuclease-NTPase systems based on oligomeric states as determined by mass photometry. All 5 models were compared in ChimeraX7 using MatchMaker^48,73^. All models for each system were relatively similar and model 0 was visualized in PyMOL and presented in the figures.

### Phage genomic DNA extraction

Phage genomic DNA was extracted as described previously^74,75^. 5 mL LB cultures supplemented with 5 mM MgCl_2_, 5 mM CaCl_2_, 1 mM MnCl_2_ were inoculated with a single *E. coli* BW25113 bacterial colony (overnight, 37°C, 230 rpm). Using the overnight culture, 20 mL fresh LB supplemented with 5 mM MgCl_2_, 5 mM CaCl_2_, and 1 mM MnCl_2_ were inoculated at an OD_600_ of 0.1 and grown at 37°C 230 rpm until an OD_600_ of 0.3 was reached. Cultures were then infected with phage at MOI of 3 and incubated until culture collapse (∼3 hours). Phage-infected cultures were centrifuged, and supernatants were passed through 0.22 µm filter. Phage supernatants were treated with DNase I and RNase A for 30 minutes at 37°C. 10 mL of phage precipitant solution (30% PEG-8000, 3 M NaCl) was added to phage supernatants and incubated at 4°C overnight. Phage particles were pelleted by spinning at 10,000×g for 10 minutes at 4°C. Resulting supernatants were decanted, and phage pellets were resuspended in 500 µL phage resuspension solution (10 mM EDTA, 5 mM MgSO_4_) and incubated with 100 µg mL^−1^ proteinase K (50°C, 30 min). Phage genomic DNA was then purified using Wizard DNA Clean-Up Kit (Promega). Briefly, phage DNA was mixed with 1 mL Wizard DNA Clean-Up Resin and added to a column. DNA was washed with 2 mL of 80% isopropanol, dried by centrifugation at max speed 2 min, and eluted with 100 µL pre-warmed 80°C distilled water. For *Bacillus* phage SPO1, phage was propagated using PY79 cells. pGEM9z was prepped by maxi-prep (Qiagen). BL21(DE3) gDNA was isolated using a DNeasy Blood and Tissue Kit (Qiagen). M13mp18 single-stranded DNA and phage Lambda DNA were purchased from NEB.

### DNA cleavage assays

Phage gDNA or plasmid DNA substrates were incubated with indicated proteins in a 20 µL cleavage reaction with 100–200 ng DNA, 10 nM – 1 µM protein, 20 mM Tris-HCl pH 7.5, 150 mM KCl, 5 mM MgCl_2_, 1 mM MnCl_2_, 1 mM DTT. Proteins were diluted in gel filtration buffer supplemented with 10% glycerol. Reactions were incubated at 37°C for 30 min followed by 100 µg mL^−1^ Proteinase K treatment at 50°C for 30 min. DNA loading buffer containing 60 mM EDTA was added to the reactions and 10 µL was loaded onto a 0.8− 1.0% TAE (40 mM Tris, 20 mM acetic acid, 1 mM EDTA) agarose gel run at 150 V for 30 min at room temperature. Gels were post-stained with TAE supplemented with 10 µg mL^−1^ ethidium bromide for 30 min, de-stained in TAE for 30 min, and imaged on a ChemiDoc MP Imaging System.

### DNA cleavage product sequencing

Nuclease-NTPase protein concentrations were optimized to generate 100–600 bp DNA fragments required for DNA sequencing. 25 nM *Pa*AbpAB, 75 nM *Bt*Hachiman, 1 µM *Ec*PD-T4-8 were individually incubated with 400 ng pGEM9Z plasmid in a 40 µL reactions with DNA cleavage buffer (20 mM Tris-HCl pH 7.5, 150 mM KCl, 5 mM MgCl_2_, 1 mM MnCl_2_, 1 mM DTT). 25 nM *Pa*AbpAB, 1 µM *Bm*Azaca, 25 nM *Bt*Hachiman, and 1 µM *Ec*PD-T4-8 were individually incubated with 400 ng phage T4 DNA in 40 µL reactions with DNA cleavage buffer. Proteins were diluted in gel filtration buffer supplemented with 10% glycerol. Reactions were incubated at 37°C for 30 min followed by 100 µg mL^−1^ Proteinase K treatment at 50°C for 30 min. 35 µL of cleavage reactions were phenol chloroform extracted, and ethanol precipitated. Briefly, 1 M Tris-HCl pH 8.0, 10% SDS, and 0.5 M EDTA pH 8.0 were added to the cleavage reaction followed by 10% SDS to reach 500 µL total volume. 500 µL of Phenol:Chlorofom:Isoamyl Alchol 25:24:1 Saturated with 10 mM Tris, pH 8.0, 1 mM EDTA (Sigma Aldrich) was added, vortexed, centrifuged, and the resulting aqueous phage was added to 50 µL 3M NaOAc and 1 µL 20 mg mL^−1^ glycogen. 1 mL of 100% ethanol was added, vortexed, and samples were stored at −20°C overnight. The following day, samples were pelleted and washed with 70% ethanol three times before resuspension in RNase-free dH_2_O.

Isolated DNA cleavage products were sent for Cut & Run sequencing at the Dana-Farber Cancer Institute Molecular Biology Core Facility (RRID:SCR_009754). Sequencing libraries were made from single stranded DNA fragments using a xGEN kit (IDT) as previously described and sequenced on a NextSeq500^54^. For bioinformatics analyses, paired-end sequencing reads were trimmed using Cutadapt to remove standard Illumina adapters, poly-A/T, and ambiguous bases. Reads were mapped to the pGEM9Z(−) plasmid (Promega) or the *Escherichia* phage T4 genome (GenBank GCA_000836945.1) using Bowtie2^55,56^. The location of the 5’ end of each read was used as the cut site and extracted from the output sam files. The 10-nucleotide sequence upstream and downstream of the cut sites were compiled and used to identify the consensus cut site using WebLogo3^76^. The lengths of the mapped reads were extracted using custom Python scripts and histogram plots were made using Prism.

**Extended Data Figure 1.**
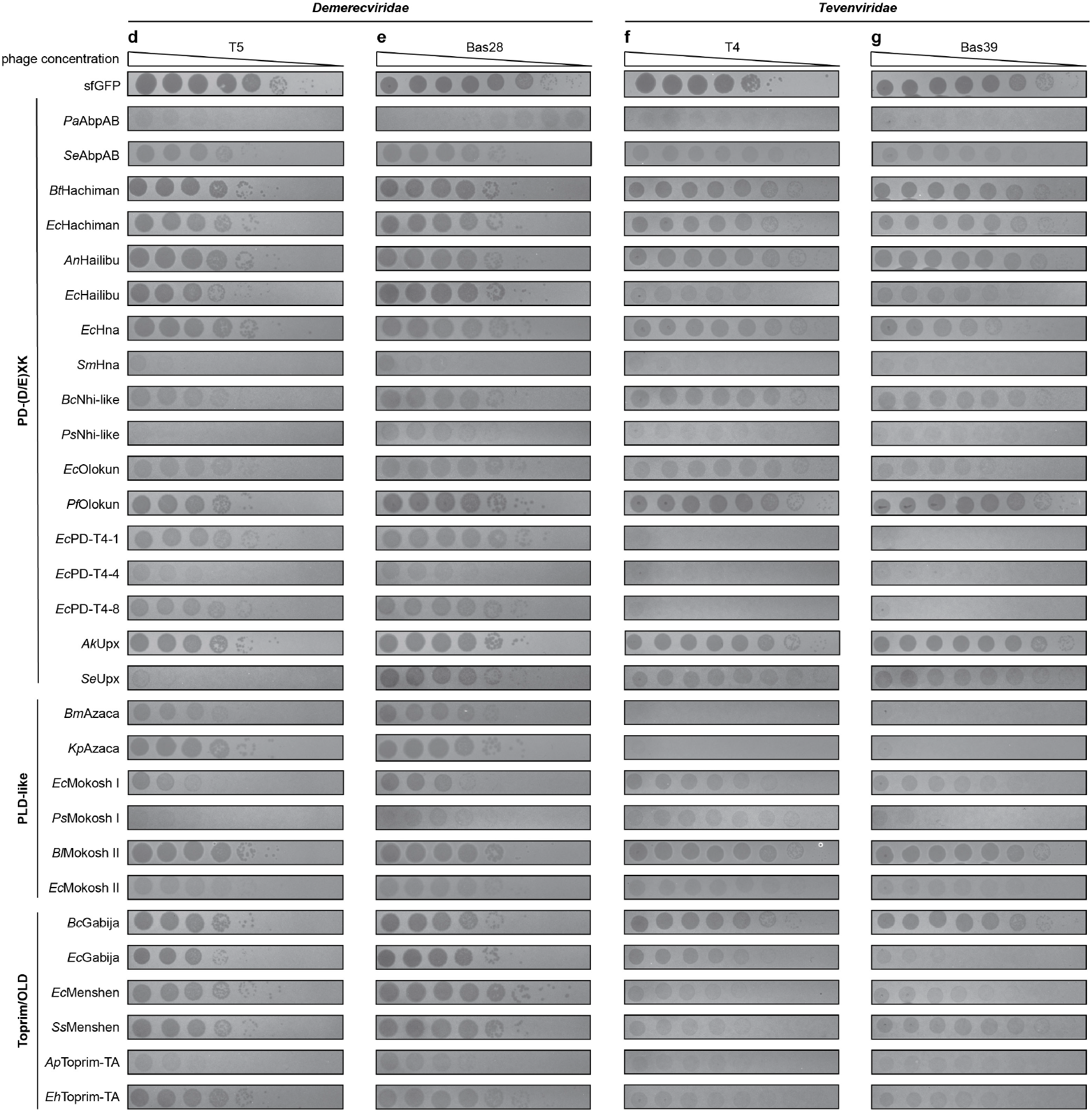
Bacteria expressing nuclease-NTPase systems are viable at low arabinose induction. Bacterial growth assay of *E. coli* expressing nuclease-NTPase operons from an arabinose inducible pBAD plasmid. Operons are displayed in alphabetical order with suppressing conditions (1% glucose) above inducing (0.02% arabinose) conditions. Data shown are representative of at least three independent experiments.

**Extended Data Figure 2.**
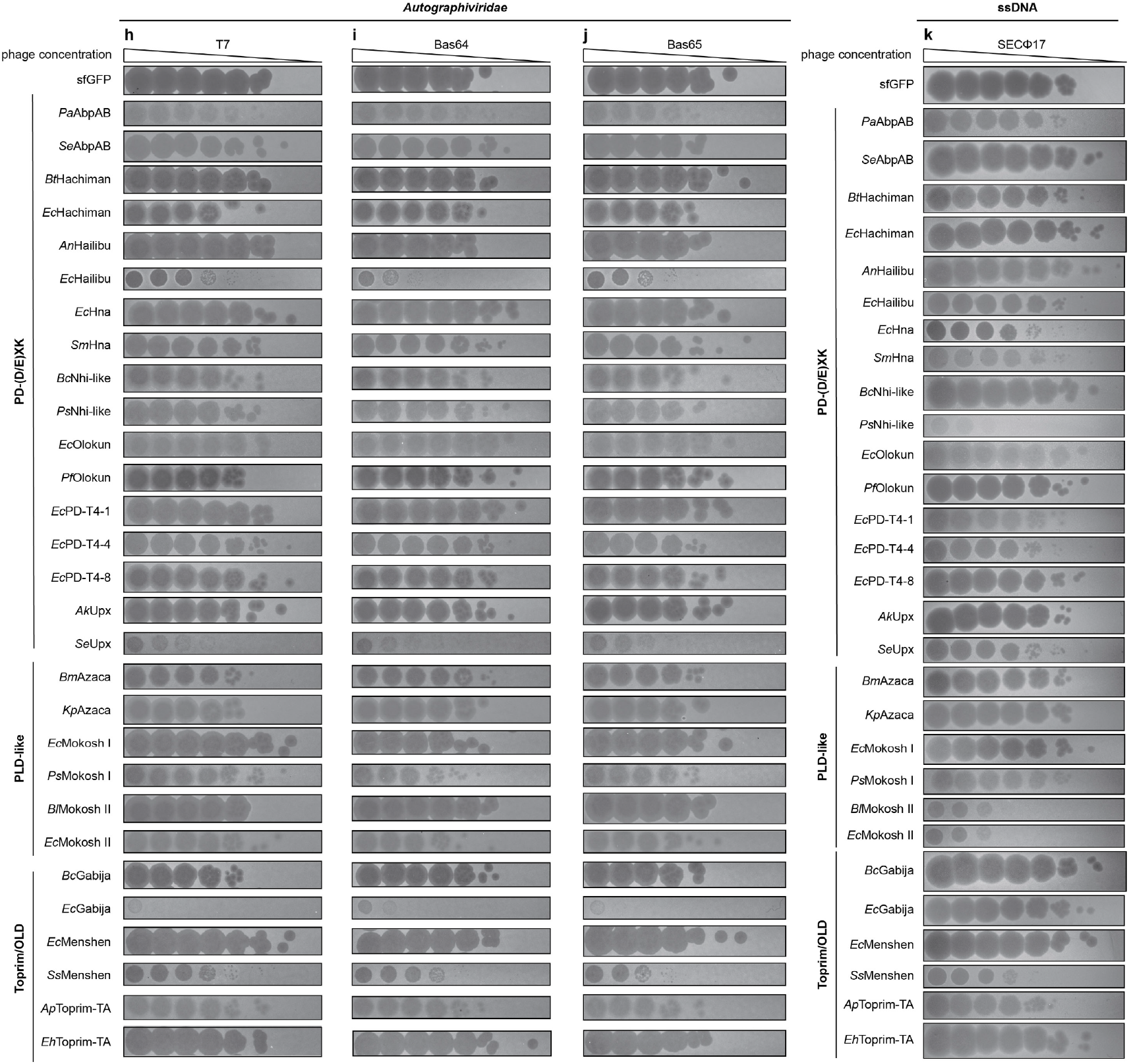
Nuclease-NTPase systems defend against diverse phages. Representative phage challenge plaque assays of bacteria expressing an sfGFP negative control or wild-type nuclease-NTPase operons and challenged with Bas17 (**a**), Bas21 (**b**), Bas25 (**c**), T5 (**d**), Bas28 (**e**), T4 (**f**), Bas39 (**g**), T7 (**h**), Bas64 (**i**), Bas65 (**j**), and SECФ17 (**k**). Data are grouped by phage and ordered by nuclease-type. Data shown are representative of three independent experiments.

**Extended Data Figure 3.**
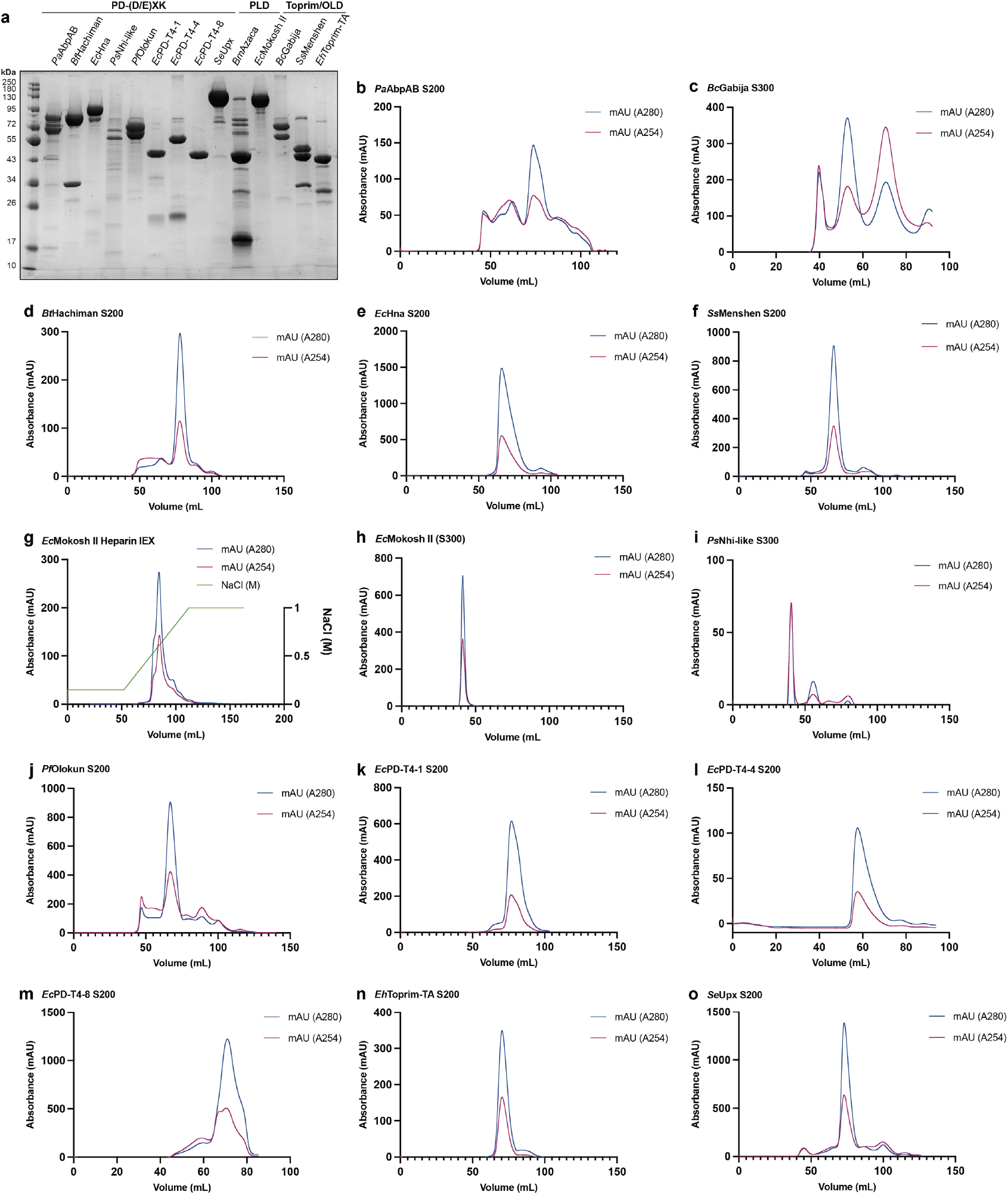
Purification of nuclease-NTPase system protein complexes. **a**, Coomassie-stained 10% SDS-PAGE gel analysis of purified nuclease-NTPase protein complexes grouped by nuclease type demonstrates that all multi-gene operons co-purify. **b–f, h–n**, Size-exclusion chromatograms (16/600 S200 or 16/600 S300) for purified nuclease-NTPase complexes. **g**, Heparin ion exchange (IEX) chromatogram for purification of *Ec*Mokosh II.

**Extended Data Figure 4.**
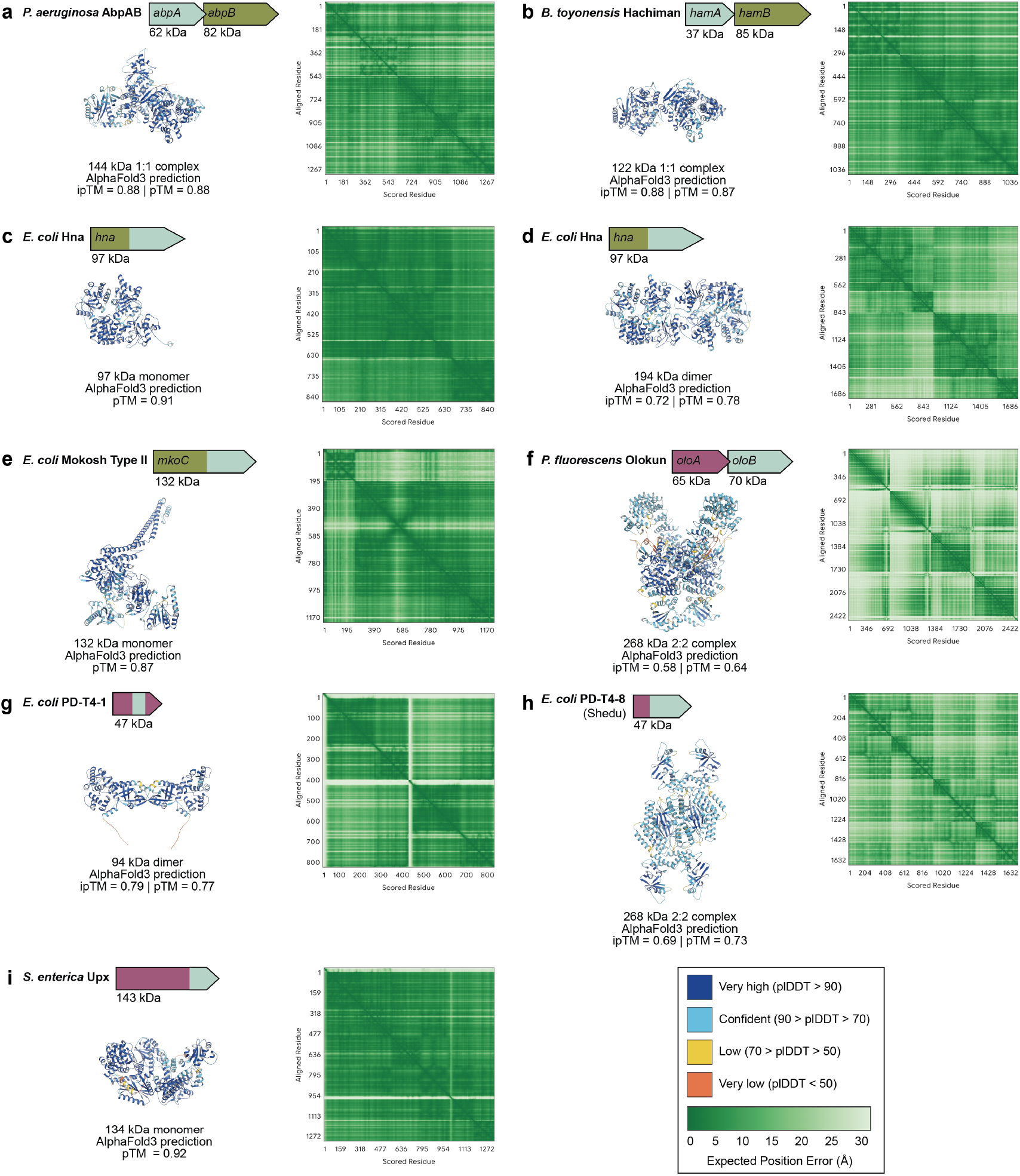
Nuclease-NTPase AlphaFold3 predictions. **a–h**, AlphaFold3 predictions of nuclease-NTPase systems colored by plDDT with expected position error graphs shown to the right of each model.

**Extended Data Figure 5.**
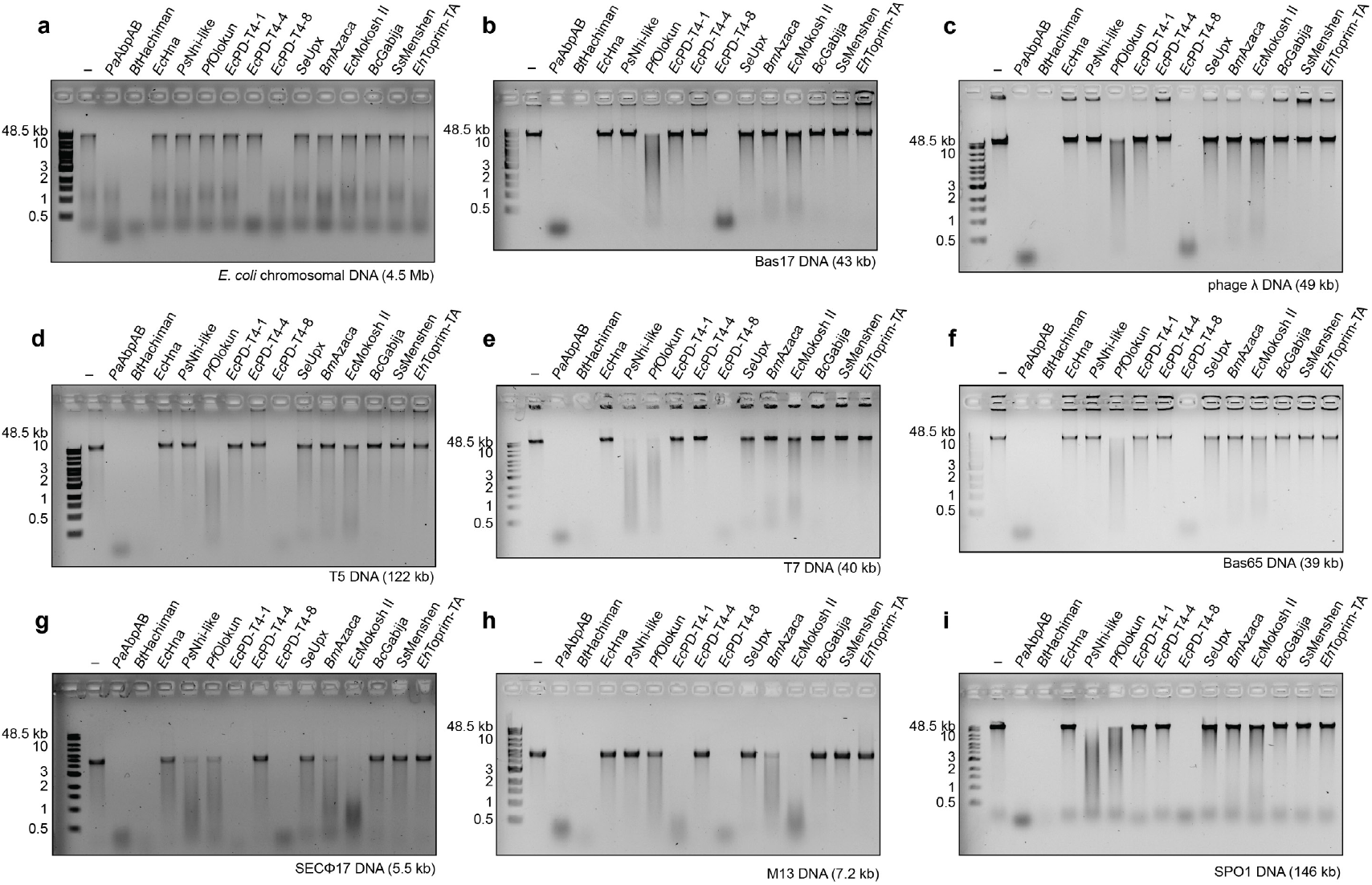
DNA cleavage assay results of all 14 purified nuclease-NTPase systems with various DNA substrates. **a–i**, DNA cleavage and agarose gel analysis of in vitro cleavage of *E. coli* chromosomal DNA (**a**), phage Bas17 DNA (**b**), phage λ DNA (**c**), phage T5 DNA (**d**), phage T7 DNA (**e**), phage Bas65 DNA (**f**), phage SECФ17 DNA (**g**), phage M13 DNA (**h**), and phage SPO1 DNA (**i**) by nuclease-NTPase systems. Data are summarized in Figure **3c** and are representative of at least 3 independent experiments.

**Extended Data Figure 6.**
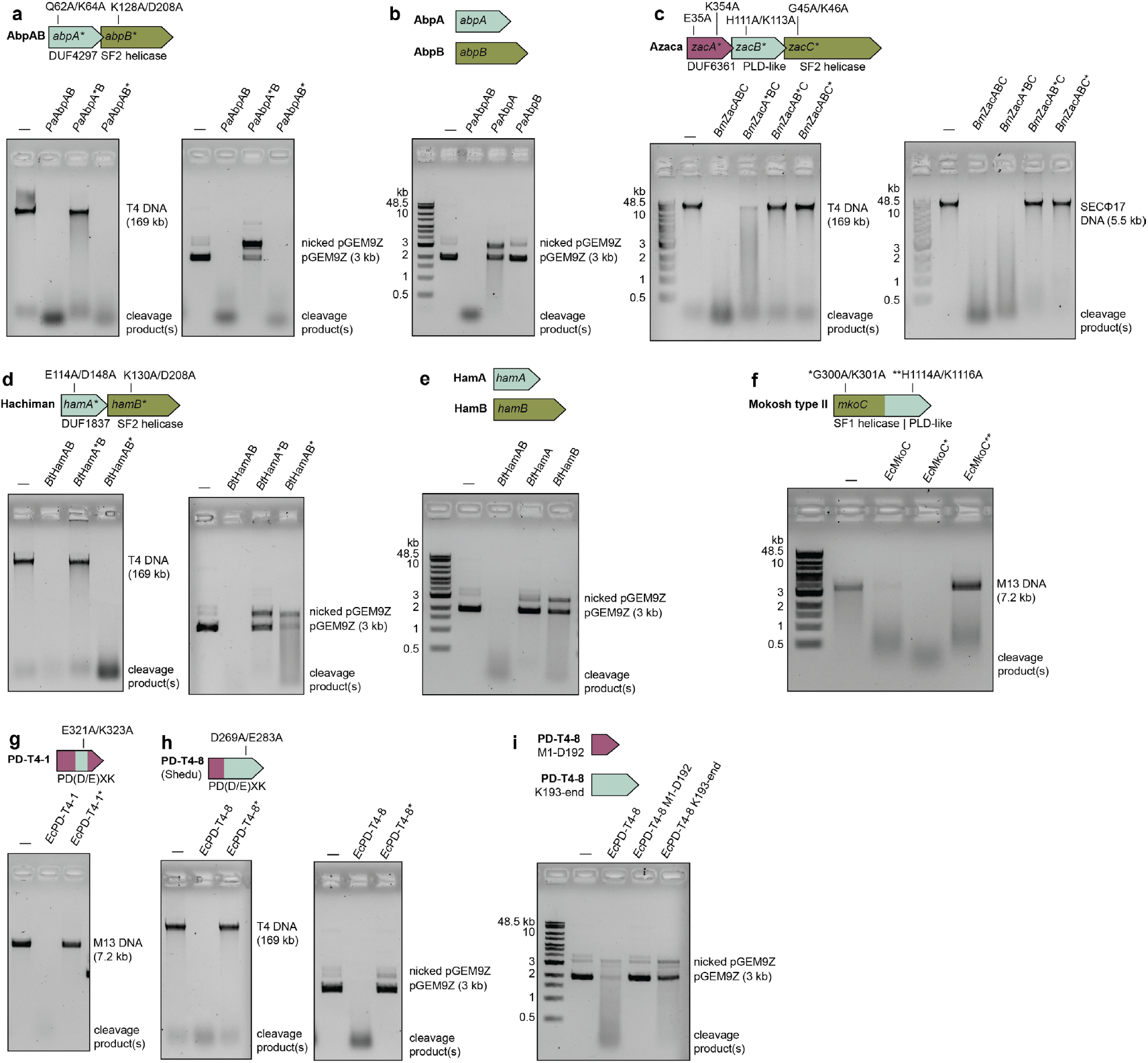
Nuclease-NTPase complexes and nuclease active site residues are required for nucleic acid cleavage. **a**, Agarose gel analysis of the ability of nuclease and NTPase catalytic mutants of *Pa*AbpAB to cleave phage T4 DNA and plasmid DNA. The wildtype NTPase active site motifs of *Pa*AbpAB are not required for cleavage. **b**, Agarose gel analysis of the ability of *Pa*AbpA or *Pa*AbpB alone to cleave plasmid DNA demonstrates the *Pa*AbpAB complex is required for cleavage. **c**, Agarose gel analysis of the ability of nuclease and NTPase catalytic mutants of *Bm*Azaca to cleave phage T4 DNA and SECФ17 DNA. Conserved residues in ZacA are dispensable for DNA cleavage, but nuclease and NTPase mutant proteins are no longer capable of cleaving phage T4 DNA. **d**, Agarose gel analysis of the ability of nuclease and NTPase catalytic mutants of *Bt*Hachiman to cleave phage T4 DNA and plasmid DNA. The wildtype NTPase active site motifs of *Bt*Hachiman are not required for cleavage. **e**, Agarose gel analysis of the ability of *Bt*HamA or *Bt*HamB alone to cleave plasmid DNA demonstrates the *Bt*HamAB complex is required for cleavage. **f**, Agarose gel analysis of the ability of nuclease and NTPase catalytic mutants of *Ec*Mokosh II to cleave M13 DNA demonstrates DNA cleavage is only inhibited when the nuclease active site is mutated. **g**, Agarose gel analysis demonstrates an *Ec*PD-T4-1 nuclease mutant is unable to cleave phage M13 DNA. **h**, Agarose gel analysis demonstrates an *Ec*PD-T4-8 nuclease mutant is unable to cleave phage T4 DNA and plasmid DNA. **i**, Agarose gel analysis of the ability of *Ec*PD-T4-8 individual subunit truncations to cleave plasmid DNA demonstrates the full-length protein is required for cleavage. Data in **a–i** are representative of at least 3 independent experiments.

**Extended Data Figure 7.**
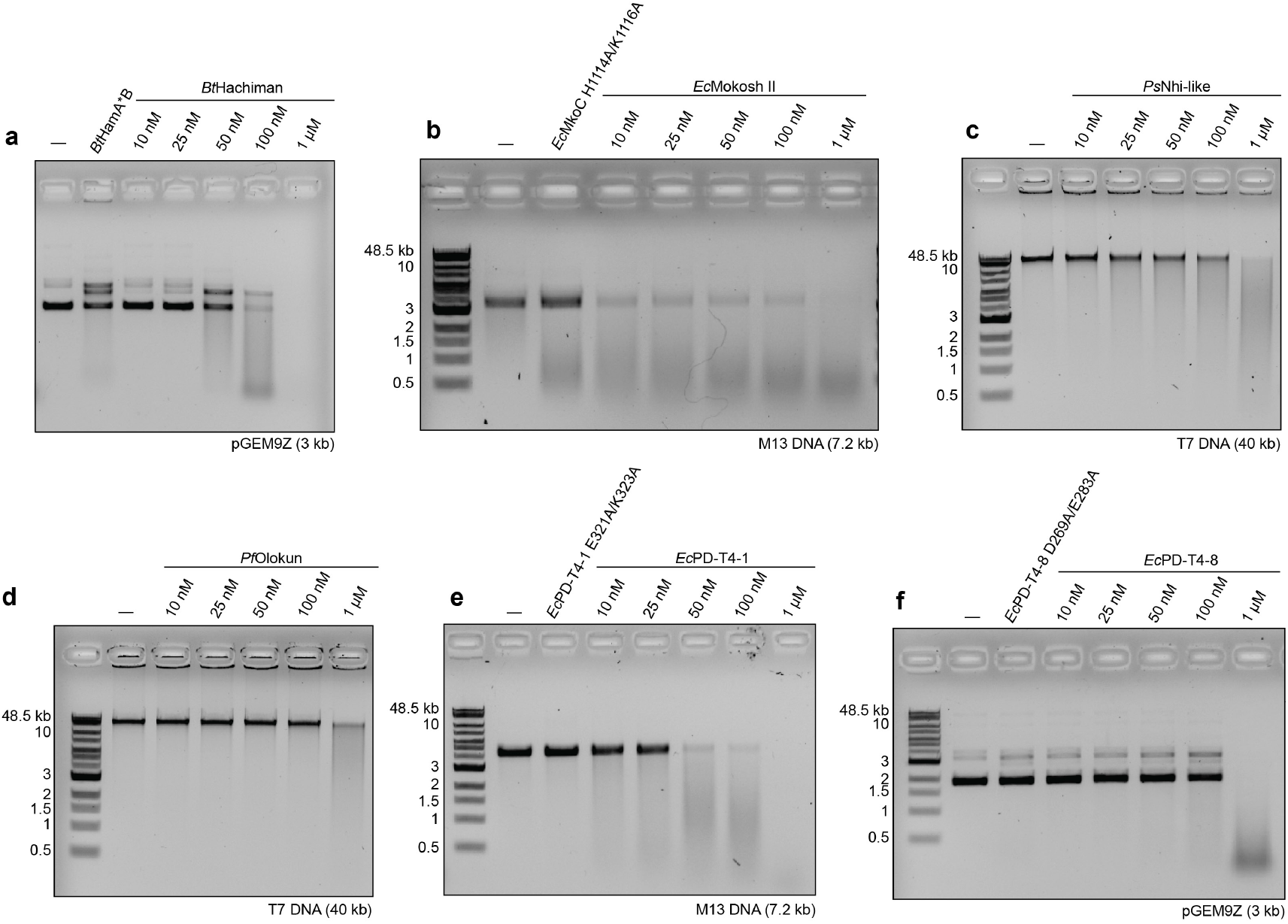
Protein titration analysis of nuclease-NTPase DNA cleavage reactions. **a–f**, DNA cleavage and agarose gel analysis of the ability of *Bt*Hachiman (**a**), *Ec*Mokosh II (**b**), *Ps*Nhi-like (**c**), *Pf*Olokun (**d**), *Ec*PD-T4-1 (**e**), and *Ec*PD-T4-8 (**f**) to cleave plasmid DNA at protein concentrations from 10 nM to 1 µM. *Pa*AbpAB, *Ec*PD-T4-1, and *Bm*Azaca cleave DNA at concentrations !50 nM. *Bt*Hachiman, *Ps*Nhi-like, *Pf*Olokun, *Ec*PD-T4-8, and *Ec*Mokosh II cleave at 1 µM. Data are summarized in Figure **4c** and are representative of at least 3 independent experiments.

**Extended Data Figure 8.**
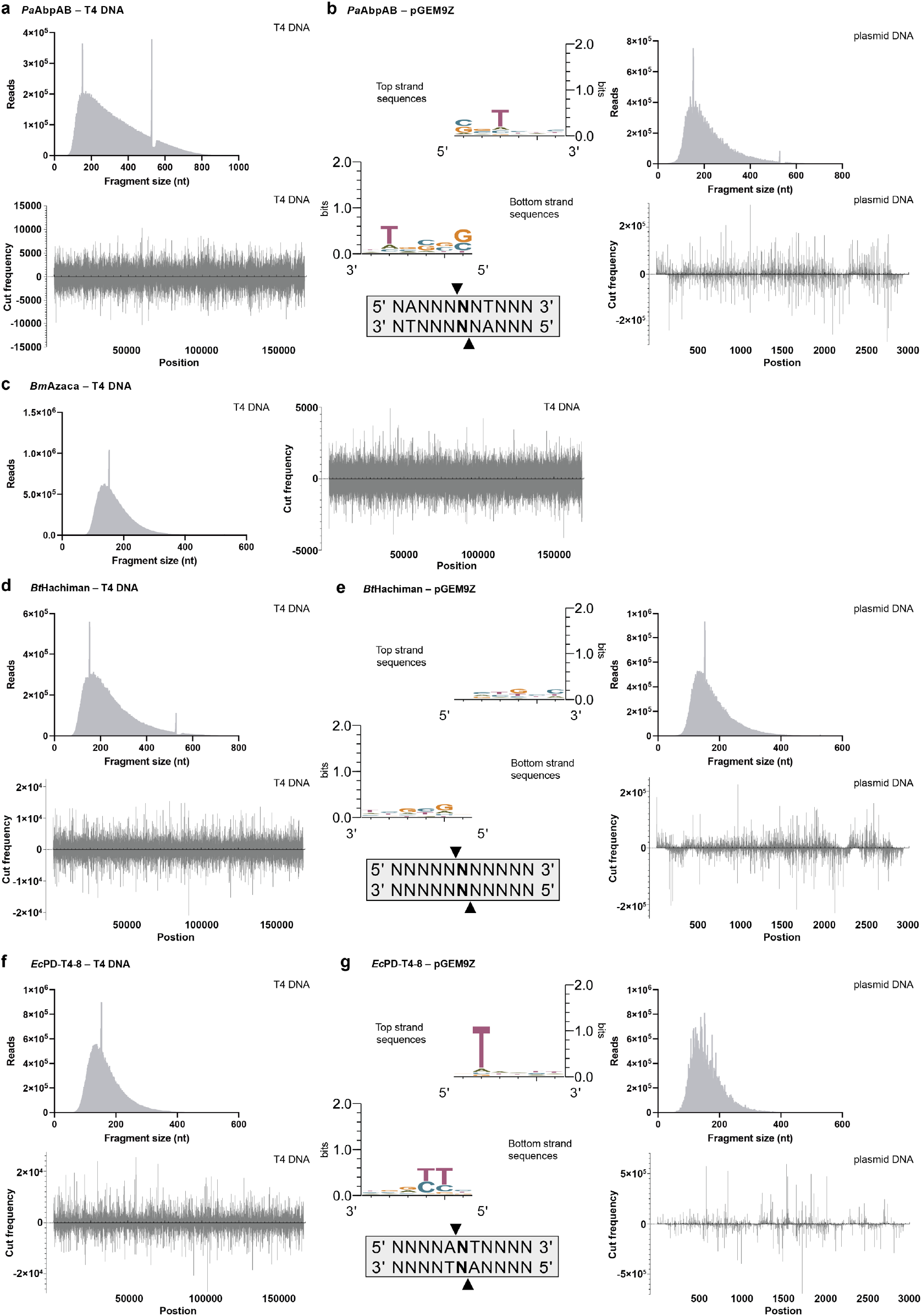
Sequencing analysis of T4 and plasmid DNA fragments incubated with nuclease-NTPase systems reveals degenerate cleavage motifs. **a, c, d, f**, Fragment size and position graphs for *Pa*AbpAB (**a**), *Bm*Azaca (**c**), *Bt*Hachiman (**d**), and *Ec*PD-T4-8 (**f**) cleavage reactions with phage T4 DNA reveals that each system cuts throughout the length of T4 DNA. **b, e, g**, WebLogos, consensus sequences, fragment size and position graphs for *Pa*AbpAB (**b**), *Bt*Hachiman (**e**), *Ec*PD-T4-8 (**g**) cleavage reactions with plasmid DNA. All three systems exhibit degenerate cleavage site motifs and cut throughout the length of plasmid DNA. All reactions were optimized such that fragment sizes would be between 150–350 bp for sequencing.

